# Default Mode Network spatial configuration varies across task domains

**DOI:** 10.1101/2021.03.17.435799

**Authors:** Lorenzo Mancuso, Sara Cavuoti-Cabanillas, Donato Liloia, Jordi Manuello, Giulia Buzi, Sergio Duca, Franco Cauda, Tommaso Costa

## Abstract

Recent developments in network neuroscience suggest reconsidering what we thought we knew about the Default Mode Network (DMN). Although this network has always been seen as unitary and associated with the resting state, a new deconstructive line of research is pointing out that the DMN could be divided into multiple subsystems supporting different functions. By now, it is well known that the DMN is not only deactivated by tasks, but also involved in affective, mnestic, and social paradigms, among others. Nonetheless, it is starting to become clear that the array of activities in which it is involved, might also be extended to more extrinsic functions. The present meta-analytic study is meant to push this boundary a bit further. The BrainMap database was searched for all experimental paradigms activating the DMN, and their activation maps were then computed. An additional map of task-induced deactivations was also created. A Multidimensional Scaling indicated that such maps could be arranged along an anatomo-psychological gradient, which goes from midline core activations, associated with the most internal functions, to the involvement of lateral cortices in more external tasks. Further investigations suggested that such extrinsic mode is especially related to reward, semantic, and emotional functions. However, an important finding was that the variability of task-induced DMN anatomic redistribution was hard to recapitulate, as none of the maps, or any linear combination of them, could represent the whole space of its dynamical reconfiguration. Altogether, our findings suggest that the DMN may be characterized by a richer functional diversity and a more spatial complexity than previously suggested.

## Introduction

Network neuroscience has partitioned the human connectome (Sporns et al., 2005) into a set of canonical networks (Damoiseaux et al., 2006; De Luca et al., 2006; Laird et al., 2011; Yeo et al., 2011). Among these, the Default Mode Network (DMN) is given special attention. At the time of writing, 6,238 published papers were returned by the PubMed search “default mode network”, while only 2,951 records for “salience network”, and 3,329 for “frontoparietal network”. At least in part, the interest for the DMN likely stems from its clinical relevance, since it is found to be altered in a wide range of diseases across psycho- and neuropathology (Mohan et al., 2016; Sha et al., 2019, 2018). However, despite the attention received by the scientific community, the function of DMN remains unclear. As a matter of fact, its elusiveness might be what makes investigating such brain system so compelling.

Initially, the DMN was outlined as a network specifically related to the resting state. Its first image comes from a meta-analysis by Shulman and colleagues (Shulman et al., 1997) about the so-called Task-Induced Deactivations (TID), depicting the areas consistently deactivated during attention-demanding tasks. Afterwards, trying to better characterize the concept of deactivation (Raichle and Snyder, 2007), Raichle and colleagues (2001) observed that the metabolism was mostly uniform across the brain during resting state. For this reason, they suggested that the brain at rest was in a state of physiological baseline (Gusnard and Raichle, 2001; Raichle et al., 2001), a *default mode* that represents a form of tonic activation for those regions commonly deactivated by tasks (Raichle et al., 2001). Finally, Greicius and colleagues (2003) proved that such default mode system was indeed a network, denoted by functional connectivity (FC) at rest. Furthermore, the DMN functional signal was reported to be negatively correlated with the signal of task-positive regions at rest (M. D. Fox et al., 2005; Uddin et al., 2009), as well as during a task and with the experimental model itself (Golland et al., 2007; Greicius and Menon, 2004; Lin et al., 2011; Newton et al., 2011). However, this *rest*-*task* distinction soon showed its limitations, in favor of an *internal*-*external* characterization of different modes of cognition (Buckner et al., 2008; Dixon et al., 2014; Fransson, 2005; Spreng, 2012).

From the early beginning of the investigations, it was observed that the DMN is not deactivated by any task, as self-referential and emotional paradigms activated it (Gusnard et al., 2001). Since those observations, many further functions were shown to be associated to this network. Other than self-referential and emotional processes (Buckner and Carroll, 2007; D’Argembeau et al., 2010; Denny et al., 2012; Engen et al., 2017; Fingelkurts et al., 2020; Fossati et al., 2003; Knyazev et al., 2020; Molnar-Szakacs and Uddin, 2013; Northoff et al., 2006; Northoff and Bermpohl, 2004; Ochsner et al., 2005, 2004; Satpute and Lindquist, 2019; Uddin et al., 2007), the DMN turned out to be related to memory and mental time-travel (Addis et al., 2007; Cabeza et al., 1997; Foster et al., 2012; Kim, 2016; Murphy et al., 2018; Rugg and Vilberg, 2013; Schacter et al., 2008, 2007; Spreng et al., 2015; Svoboda et al., 2006; Yang et al., 2013), mental simulation and scene construction (Gerlach et al., 2011; Hassabis et al., 2007; Hassabis and Maguire, 2007; Spreng and Grady, 2010), theory of mind (ToM) and social cognition (Amft et al., 2015; Mar, 2011; Mars et al., 2012; Mwilambwe-Tshilobo and Spreng, 2021; Rilling et al., 2004; Ruby and Decety, 2004; Saxe and Kanwisher, 2003; Saxe and Powell, 2006; Spreng and Andrews-Hanna, 2015), moral judgment (Bzdok et al., 2012; Greene et al., 2001; Harrison et al., 2008; Pujol et al., 2008), semantic processing (Binder et al., 1999, 2009; Chiou et al., 2020; Evans et al., 2020; Lanzoni et al., 2020), and reward mechanisms (Lopez-Persem et al., 2020; Martins et al., 2021; Xue et al., 2009). However, most of these psychological functions can still be somewhat associated with the resting state (Wen et al., 2020). In fact, the resting mind has not to be considered as idle (Raichle, 2009), as it is continuously involved in activities collectively known as *mind-wandering* (Fox et al., 2015; Giambra, 1989; Mason et al., 2007; Seli et al., 2016; Spiers and Maguire, 2006). During mind-wandering, the brain is expected to engage in DMN-related functions such as remembering the past, imagining the future, thinking about others, and displacing the self in imaginary situations (Andreasen et al., 1995; Andrews-Hanna, 2012; Andrews-Hanna et al., 2014; Christoff et al., 2016). When some of these activities intrude into the execution of an attentional task as *stimulus-independent thoughts*, the task performance may be affected by errors (Kam and Handy, 2013; Prado and Weissman, 2011; Smallwood et al., 2008; Sonuga-Barke and Castellanos, 2007). Likewise, expected task-related activations may be found disrupted, and DMN areas activated (Eichele et al., 2008; Kam et al., 2011; Li et al., 2007; Mason et al., 2007; Weissman et al., 2006). Still, these observations are still consistent with a *rest*-*task* dichotomy, and do not really suggest a proactive role for the DMN. In contrast, it has been proposed that, during task, DMN-driven *stimulus-oriented thoughts* may possibly appear (Mantini and Vanduffel, 2013; Sormaz et al., 2018) with the purpose of supporting task performance (Gilbert et al., 2007). As a matter of fact, the DMN has been implicated in problem-solving and creativity (Abraham et al., 2012; Benedek et al., 2014; Ellamil et al., 2012; Gerlach et al., 2011; Huo et al., 2020; Jung et al., 2013; Kounios et al., 2006; Marron et al., 2018; Mayseless et al., 2015), suggesting that its activity can also be engaged in external demands. Indeed, DMN activations can enhance cognitive control during extrinsic tasks requiring internal mentation (Spreng et al., 2014), possibly in coordination with frontoparietal control systems (Cocchi et al., 2013; Gerlach et al., 2014; Spreng et al., 2010). Furthermore, the anteromedial prefrontal cortex (amPFC), node of the DMN, was found to be activated in monitoring the external environment, contributing to faster reaction times (Gilbert et al., 2006, 2005). It also seems that the DMN is recruited in switching tasks, in the case of a demanding shift from a cognitive context to a different one (Crittenden et al., 2015). Moreover, the DMN was also found to be activated when subjects have to automatically apply learned rules (Vatansever et al., 2017).

The critical role of DMN for task execution is highlighted not only by activation studies, but also by functional connectivity investigations that questioned the supposed DMN anticorrelation with task execution and task-positive regions. It has been shown that nodes of the DMN are positively correlated with task-positive areas during acquisition and retrieval phases of a working memory task (Piccoli et al., 2015), as well as during the preparation phase (Koshino et al., 2014, 2011). Also, during such tasks, the connectivity within the DMN was found to be correlated with behavioral performance (Hampson et al., 2006). Moreover, the sign and the strength of the correlations between DMN and task-positive regions remarkably varies between nodes of these networks, and according to the tasks and the different resting state time epochs (Bluhm et al., 2011; Chang and Glover, 2010; Denkova et al., 2019; Dixon et al., 2017; Elton and Gao, 2015; Leech et al., 2011). At the subject level, interactions between DMN and attentional and control networks were found during a semantic memory retrieval task (Fornito et al., 2012). In sum, there is considerable evidence that the DMN functionality is crucial, not only for internal mind-wandering, but also for the execution of extrinsic activities.

Just as its role in human cognition, the anatomical and topological representation of the DMN has proven to be puzzling. From a theoretical point of view, the DMN could be described as a whole and unfractioned network, with hub nodes in the posterior cingulate cortex (PCC) and medial prefrontal cortex (mPFC) and more peripheral nodes in the medial and lateral temporal lobe, angular gyrus (AG), dorsolateral prefrontal cortex (dlPFC), and inferior frontal gyrus (IFG) (Buckner et al., 2008; Yeo et al., 2011). Nonetheless, it is widely acknowledged that it can be further divided into subnetworks (Abou-Elseoud et al., 2010; Abou Elseoud et al., 2011; Ray et al., 2013; Shirer et al., 2012; Yeo et al., 2011). In this regard, Andrews-Hanna and colleagues (2010) reported that the subdivisions of such network might have different functions. In particular, they identified two subsystems structured around a midline core: the former, formed by the PCC was thought to be involved in self-referential processes, and the latter, composed by mPFC, has considered to be related to future-oriented thoughts (Andrews-Hanna et al., 2010). What is more, the midline core itself might be a fractionated structure. In fact, by analyzing single-subject data with minimal spatial smoothing, Braga and colleagues (Braga et al., 2019; Braga and Buckner, 2017; DiNicola et al., 2020) have recently found that the DMN seems to be composed of two parallel and interdigitated networks, also interleaved within the PCC and mPFC. The two subsystems were found to be related to different roles: one to social cognition and the other one to mnestic functions (DiNicola et al., 2020). Similarly, Wang and colleagues (2020) parcellated the DMN nodes into different parts, each one associated with a specific functional profile. Likewise, Gordon and colleagues (2020) were able to divide the individuals’ DMN into nine subnetworks showing differential task engagement. Thus, the most recent developments in the research of the DMN are indicating that such network, far from being a monolithic entity, consists of multiple systems with intersecting functions and anatomies (Buckner and DiNicola, 2019).

The present study aims to delve into this matter, using a coordinate-based meta-analytical methodology to investigate the functions related to the activity of the DMN regions and the spatial variability associated with them. The use of a meta-analytical approach allows to overcome the heterogeneity of results, typical issue of neuroimaging experiments (Botvinik-Nezer et al., 2020). After all, the DMN research has a long meta-analytical tradition. In fact, the first images of the network come from meta-analyses of TID (Mazoyer et al., 2001; Shulman et al., 1997), subsequently confirmed by an Activation Likelihood Estimation (ALE) meta-analysis by Laird and colleagues (Laird et al., 2009). Another ALE study (Schilbach et al., 2012) showed that the areas of TID, social cognition, and emotional processing converged on PCC and mPFC. Similarly, an ALE meta-analysis from Spreng and colleagues (2009) noted a correspondence between autobiographical memory, spatial navigation, theory of mind activations, and TID. It could be said that these two latter studies used a meta-analytic approach to put forward a consistent and comprehensive view of DMN functions. On the contrary, the current study wants to highlight the functional variety of the DMN and the resulting modulations of its spatial configuration.

In order to make these assessments, we performed a Paradigm Analysis (Lancaster et al., 2012), capitalizing on the BrainMap database and on its taxonomy of behavioral ontologies (P. T. Fox et al., 2005; Fox and Lancaster, 2002; Laird et al., 2005). This allowed us to identify the task categories significantly associated with the network in a data-driven fashion. Activation coordinates of such paradigms were then obtained from the same database, and used to perform an ALE meta-analysis for each one of them. As indicated by Raichle and colleagues (Raichle et al., 2001), TID correspond to rest tonic activations, and it has been suggested that the rest should be seen just as another active state (Buckner et al., 2013). Thus, a TID ALE map was calculated to represent the DMN configuration during resting state. This also constitute a replication of Laird et al. (Laird et al., 2009) with a larger database and updated algorithms. The resulting set of maps underwent a series of analyses, namely, Multidimensional Scaling (MDS), Principal Component Analysis (PCA), and ICA. These were to explain the DMN task-based variability in the form of axes along which the network arranges itself to meet external demands. We expected that different operative domains related to the DMN would recruit distinctive sets of areas, including some regions typically assigned to other resting state networks (RSNs). Given the wide range of functions implicated with the mPFC (Delgado et al., 2016; Hiser and Koenigs, 2018; Lieberman et al., 2019; Schneider and Koenigs, 2017; Toro-Serey et al., 2020), we anticipated that this region would express a large variability in its activations, possibly organized in a rostral-caudal arc revolving around the callosal genu (Amodio and Frith, 2006). The PCC might show some internal differentiation as well (Leech et al., 2011).

## Methods

### Paradigm Analysis

The Paradigm Analysis (Lancaster et al., 2012), as implemented in the dedicated plugin for Mango (http://ric.uthscsa.edu/mango/) (Lancaster et al., 2011, 2010), was performed in order to get the profile of involvement of the DMN with different fMRI paradigms. However, numerous and different functional parcellations of the human brain exist, and there are no methodological criteria or gold standard to prefer one to the others (Arslan et al., 2018; Eickhoff et al., 2015). Therefore, to maximize the representativity of our results, three different masks of the DMN obtained with different methodologies were fed to the Paradigm Analysis. Only the paradigms that were found significant in at least two out of the three masks, were taken in consideration in the further analyses. The first one was extracted from the 7 Network parcellation by Yeo and colleagues (Yeo et al., 2011), which was produced with a clustering algorithm. The second one was derived from the ICA by Shirer and colleagues (Shirer et al., 2012), merging the originally split ventral and dorsal components of the DMN. Lastly, we selected the DMN from the CAREN atlas by Doucet and colleagues (Doucet et al., 2019), which was produced as the consensus between six different parcellations. Since the Paradigm Analysis tool works in Talairach space, the three masks were registered to Talairach using FLIRT (Jenkinson et al., 2002).

### Activation Likelihood Estimation and Fail-Safe analysis

In order to trace the studies related to the 8 significant paradigms identified by such consensus approach (see Results), the software Sleuth has then been used to search the BrainMap funtional database (P. T. Fox et al., 2005; Fox and Lancaster, 2002; Laird et al., 2005) eight times, composing the queries as follows:

> *[Experiment Context is Normal Mapping] AND [Experiment Activation is Activations Only] AND [Experiment Paradigm Class is …]*

with the latter field completed according to the Paradigm Analysis results. Furthermore, to obtain the TID, we replicated the search used by Laird and colleagues (Laird et al., 2009):

> *[Experiment Context is Normal Mapping] AND [Experiment Activation is Deactivations Only] AND [Experiment Control is Low Level]*

Coordinates were exported by Sleuth software in Talairach space. To minimize the within group effects and ensure independence between the observations (Müller et al., 2018; Turkeltaub et al., 2012), the experimental contrasts calculated on the same group of subjects were merged in a single set of foci, using the pertaining option in the tool.

Each one of the resulting lists of coordinates was then fed to GingerALE 3.0.2 (Eickhoff et al., 2009, 2012; Turkeltaub et al., 2002) to calculate its ALE map. A family wise error (FWE) correction for multiple comparisons was adopted (Eickhoff et al., 2012), with cluster-level inference of *p* < 0.05 and a cluster-forming threshold of *p* < 0.001.

To take into account the file-drawer effect, a Fail-Safe procedure was implemented (Acar et al., 2018). Briefly, a given number of experiments was added to each of the nine datasets to simulate unpublished results. The number of foci and subjects of those experiments were such to match the distribution they had in the original data. Samartsidis and colleagues (Samartsidis et al., 2020) estimated that the fail-drawer effect of the BrainMap database amounts to the 6%. Thus, we decided to perform a series of Fail-Safe analyses adding the 6% and the 60% of random experiment to each dataset, so as to evaluate the robustness of our results.

### Data analysis

The subsequent analyses were carried out in Python, using the NiBabel 3.2.1 package (Brett et al., 2020) to access the NIfTI file format, the NumPy library (Harris et al., 2020) to calculate the Pearson correlation coefficients and sci-kit learn 0.24.1 (Pedregosa et al., 2011) to compute MDS, PCA and ICA.

To start with, the 9 ALE maps were vectorized using a Talairach standard as brain mask with 2 mm^3^ voxel size (https://www.brainmap.org/ale/colin_tlrc_2x2x2.nii.gz), to obtain a *voxel* × *map* matrix. To perform an MDS, the Pearson correlation matrix was derived first. Then, the distance measure of the dissimilarity matrix was calculated as the 1-*correlation*, as in Kriegeskorte and colleagues (2008). The *voxel* × *map* matrix was also used as input for a PCA to obtain both the principal component (PC) loadings (map coefficients for each component) and scores (voxel projections on the component). The scores were then plotted on the Talairach standard to obtain PC maps. Finally, the same *voxel* × *map* matrix was fed to the scikit-learn FastICA algorithm (Hyvarinen, 1999; Hyvärinen and Oja, 1997). In doing so, we obtained each component coefficient, or loading, from the unmixing matrix, and the independent component (IC) voxel-wise maps.

## Results

### Paradigm analyses and Activation Likelihood Estimations

The Paradigm Analysis results obtained from the three selected DMN masks (Doucet et al., 2019; Shirer et al., 2012; Yeo et al., 2011) are presented in Table 1. Eight paradigms were found to be significant in at least 2 out of the 3 analyses: ToM, Semantic Monitor/Discrimination, Episodic Recall, Emotion Induction, Self-Reflection, Deception, Imagined Object/Scenes, and Reward. Most of these paradigms are related to social, mnestic, or other internal mentation functions typically associated with the DMN (Buckner et al., 2008). To the best of our knowledge, deception tasks were not previously related to the network, although the social nature of such paradigms likely justifies this result. Similarly, we are not aware of many explicit links between reward functions and DMN in the literature (Martins et al., 2021), albeit there is strong evidence to associate the mPFC to such mechanisms (Hiser and Koenigs, 2018; Lieberman et al., 2019; Schneider and Koenigs, 2017; Xue et al., 2009). Interestingly, reasoning and problem-solving paradigms had a significant effect on the CAREN mask (Doucet et al., 2019), suggesting that the DMN might then play a role outside of what is considered internal mentation in the strictest sense.

**Table 1:**
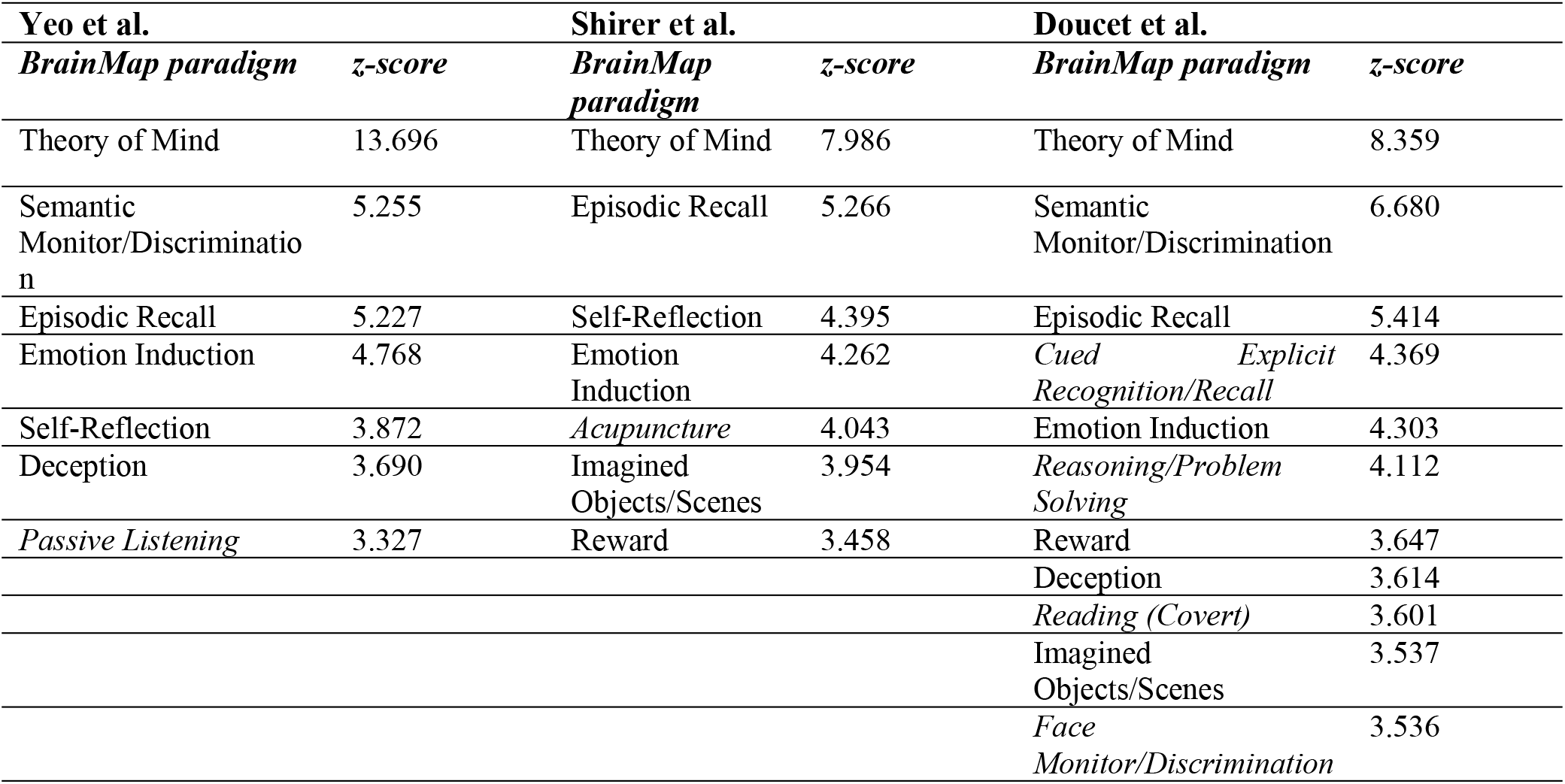
BrainMap paradigms found to be significantly associated with the DMN masks obtained by the 7 networks atlas by Yeo et al. (2011), the Independent Component Analysis by Shirer et al. (2012), and the CAREN atlas by Doucet et al. (2019). The paradigms excluded from further analysis are written in italics.

Details about the results from Sleuth searches for the 8 queries associated with each significant paradigm are presented in Table 2 and in the Prisma Flow chart in Supplementary Fig. S1. We point out that we found a limited number of experiments for the Self-Reflection condition. According to Eickhoff and colleagues (Eickhoff et al., 2016; Müller et al., 2018), at least 17 experiments should be gathered in order to perform a statistically sound ALE. Although the query returned 28 experiments, after merging them according to groups of subjects (Müller et al., 2018), they became 7 foci groupings. To rule out possible bias induced by the inclusion of this underpowered domain, the subsequent analyses were repeated with and without the related ALE map. Since excluding this condition did not significantly influence the outcome of, we decided to keep it in the dataset.

**Table 2:**
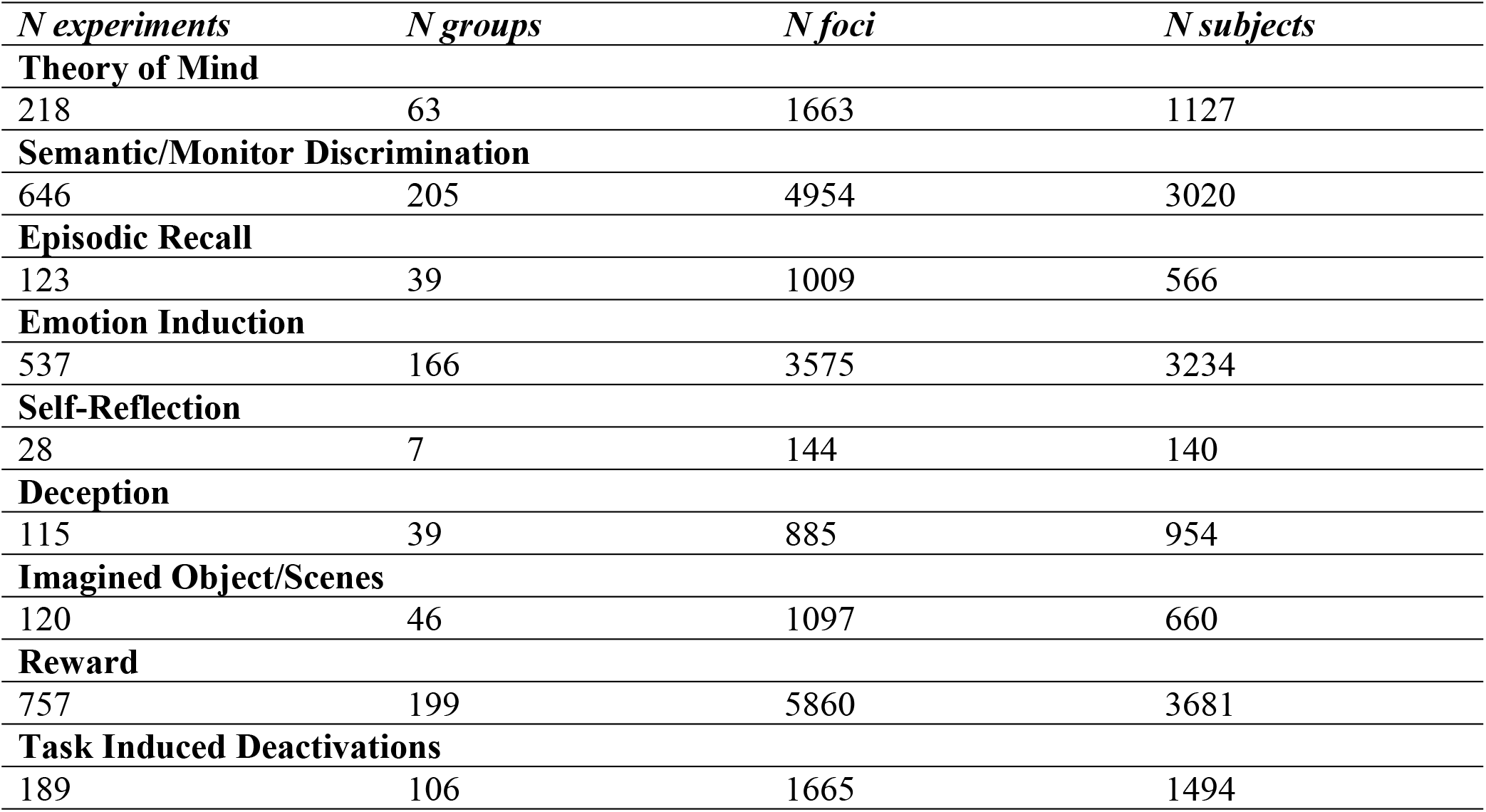
Details about the result of Sleuth queries.

We also computed a TID meta-analysis, representing rest. The ALE results are presented in Fig. 1. The TID ALE map replicates the one by Laird and colleagues (Laird et al., 2009), with the exception of the mPFC cluster, that we found in a more dorsal position.

**Figure 1:**
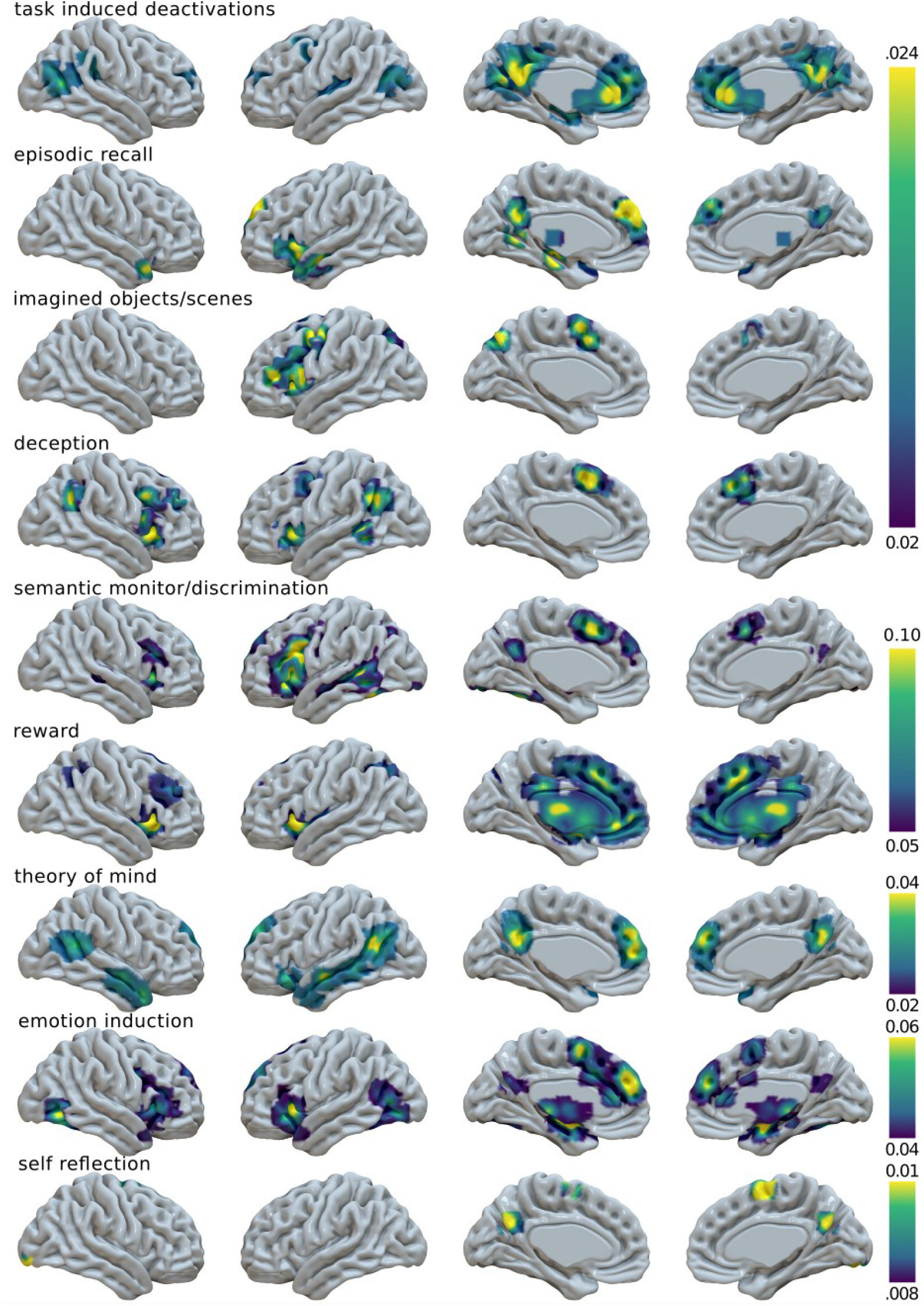
Surface mapping of the 9 Activation Likelihood Estimation maps.

As for the other maps, Episodic Recall activates the left insula along with DMN areas. Imagining objects and scenes activates the left hemisphere in the precuneus, lateral prefrontal cortex (lPFC) including IFG, and supplementary motor area (SMA). Deception tasks activate the bilateral insula, other than AG and SMA, in accordance to Farah and colleagues (Farah et al., 2014). Semantic monitoring and discrimination activate the left lPFC including IFG, but also the SMA, the left lateral and medial temporal lobe, and the PCC. Reward activates the basal ganglia (BG), the thalamus, the whole mPFC, and the bilateral insula. ToM activates the temporal and temporoparietal cortices along with PCC and dmPFC, especially on the left. Inducing emotion activates the left insula and the bilateral occipito-temporal cortices, along with typical DMN nodes. The Self-Reflection map involves PCC, the right SMA, and an occipital cluster in the fusiform gyrus. The maps obtained by the Fail-safe analyses, presented in Supplementary Fig. S2, show that our ALE maps remain substantially unchanged when accounting for the file-drawer effect.

In summary, few maps matched the prototypical representation of the DMN. Some of them showed either weak activation or no activation at all in the midline core, and a strong expression of lateral areas of the network such as AG, IFG, and middle temporal gyrus. In addition, the insula and SMA/dorsal ACC, hubs of the salience network (SN), were often present. Rather than considering these as spurious findings, we see them as an indication that, when the brain is engaged by external demands, multiple networks including DMN nodes would emerge. Although relying on intrinsic brain topology, such recruitment would be not strictly constrained by it (Cole et al., 2014; Krienen et al., 2014). Thus, it might involve a flexible shift in brain hubness (Cole et al., 2013; Fransson and Thompson, 2020) and a remodulation of cooperative and competitive long-range connectivity patterns (Dixon et al., 2017; Fornito et al., 2012; Piccoli et al., 2015).

### Multidimensional Scaling

The 1-*correlation* dissimilarity matrix (Kriegeskorte et al., 2008) between maps and the resulting 2- and 3-dimensional MDS can be seen in Fig. 2. The MDS solutions computed excluding the Self-Reflection condition are presented in Supplementary Fig. 3. In both analyses, the 2-dimensional MDS solution seems to suggest a first axis standing for medial-lateral, or a core-lPFC, spatial representation. The maps found on one side of the axis (e.g. TID, ToM, Episodic Recall) show activations in the midline DMN core, while those found on the other side (e.g. Deception, Imagined Objects/Scenes, Semantic Monitor/Discrimination) display a weaker involvement of such regions, especially PCC. Conversely, the latter have significant clusters in the lPFC and insula, and the midline activations are especially located in the SMA. The maps activating the midline core are not only those presenting more similarity with its stereotypical image, but also those whose functions were more commonly associated with the DMN (Buckner et al., 2008). As for the opposite side, these maps are related to tasks more rarely associated with the network, and they bear a similarity with the spatial distribution of SN, rather than the one of DMN. Moreover, they are reminiscent of the semantic regions (i.e., SemN) found by Chiou and colleagues (2020) as belonging to a more outward-leaning DMN subsystem (see also Evans at al. 2020). Hence, this anatomical midline-lateral axis could also be seen as a psychological internal/external dimension. As for the second axis, the distribution of the maps may suggest a dorsal-ventral labeling. On one side, activations are more focused in SMA, PCC, or dorsal frontal and parietal cortices. On the other side, there are insular, temporopolar, and medial temporal areas.

**Figure 2:**
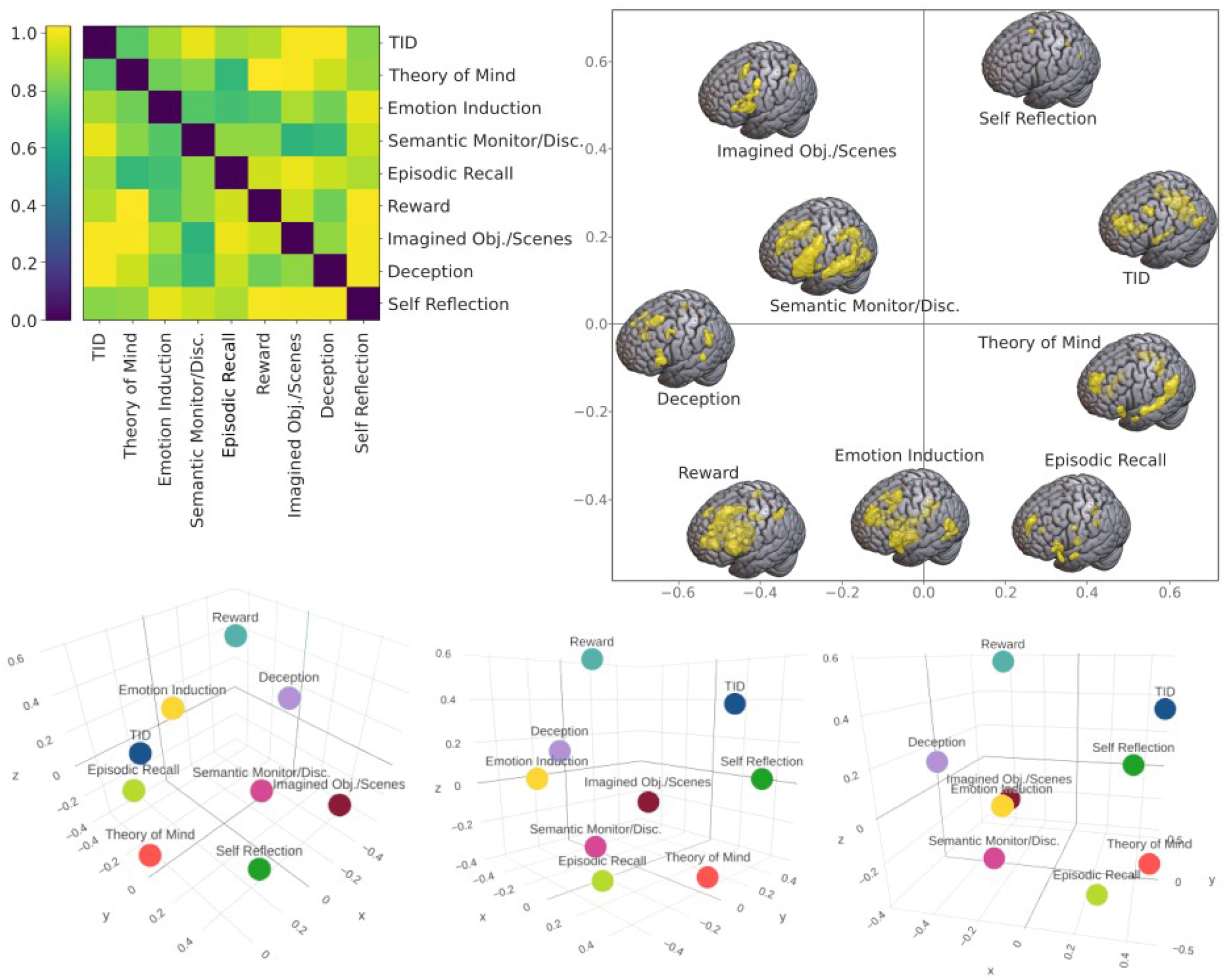
Multidimensional Scaling (MDS) of the 9 Activation Likelihood Estimation (ALE) maps. Top Left: 1-correlation distance matrix of the 9 ALE maps. Top Right: MDS 2-dimensional solution. The surface maps are centered on the MDS coordinates. Bottom: MDS 3-dimensional solution, seen from different perspectives.

Lastly, it is interesting to observe from the MDS graph that the maps seem to be placed in a circle. In other words, they look equidistant from each other and from a hypothetical center point. Yet, this central point remains empty, suggesting that none of the maps, not even the TID one, provides a good representation of all the systems implicated by the activity of DMN nodes. This intuition holds also at a higher dimensionality, as shown in the 3-D solution (Fig. 2). Even if the third axis is harder to interpret, it can be seen as the maps seem to be placed on the surface of a sphere.

### Principal Component Analysis

Given the lack of common ground between different DMN expressions, we performed a PCA to summarize the inter-paradigm similarities. However, the sphericity implied by MDS raises doubts on PCA ability to reduce our data dimensionality. Even if the high significance of a Bartlett’s test (*p* < 0.001) seems to suggest otherwise, the Kaiser-Meyer-Olkin index (KMO = 0.61) confirms a sufficient but mediocre relation between the maps.

Nonetheless, the first component (PC1) explains 51.8% of the between-map variance. The voxel-wise scores on PC1 indicate an involvement of the mPFC comprising SMA, two separated caudal and rostral PCC clusters, thalamus and BG (caudate head and anterior lenticular nucleus), anterior insula (AI), IFG, and a cluster in the lateral Brodmann area (BA) 7. This component loads especially on the Reward map, and, to a minor extent, on the Semantic Monitor/Discrimination and Emotion Induction maps.

The second component (PC2) explains 30.5% of the total variance and loads especially on the Semantic Monitor/Discrimination map. It has a weaker representation in mPFC with positive scores in SMA and dmPFC (with a leftward lateralization), but negative scores in bilateral ACC. There is a positive cluster in the caudal PCC, while the more rostral one, along with thalamus and anterior BG, scores negatively. The left lPFC (including IFG), AI, and most of the premotor cortex, form a large positive cluster. On the other hand, there is less recruitment of the right frontal cortices. Similarly, the left temporal cluster is wider than the right one, including the superior temporal sulcus up to the AG, and extending to the occipito-temporal cortex, and the fusiform gyrus. The bilateral occipital poles also score positively on the component.

The anatomical representations of the first two components are remarkably similar, mostly differing because of the involvement of BG, ACC and vmPFC, and a more anterior segment of PCC, included in PC1, but having a negative score in PC2. Also, they both show a partial left lateralization, reflecting the higher involvement of the left hemisphere in many of the investigated tasks (Fig. 3). Their spatial resemblance seems to indicate that, despite being orthogonal, these two components may be related to a similar phenomenon. In fact, we note that they both resemble the SemN (Chiou et al., 2020; Evans et al., 2020; Noonan et al., 2013). Furthermore, they remind the transitional module serving an integrative function with the frontoparietal task-positive system observed by Fornito and colleagues (2012), as well as with the Overlapping Community 6 found by Najafi and colleagues (2016). Consequently, we suspect that they both serve some form of integration between the DMN core and the task-positive areas for the execution of more external tasks.

**Figure 3:**
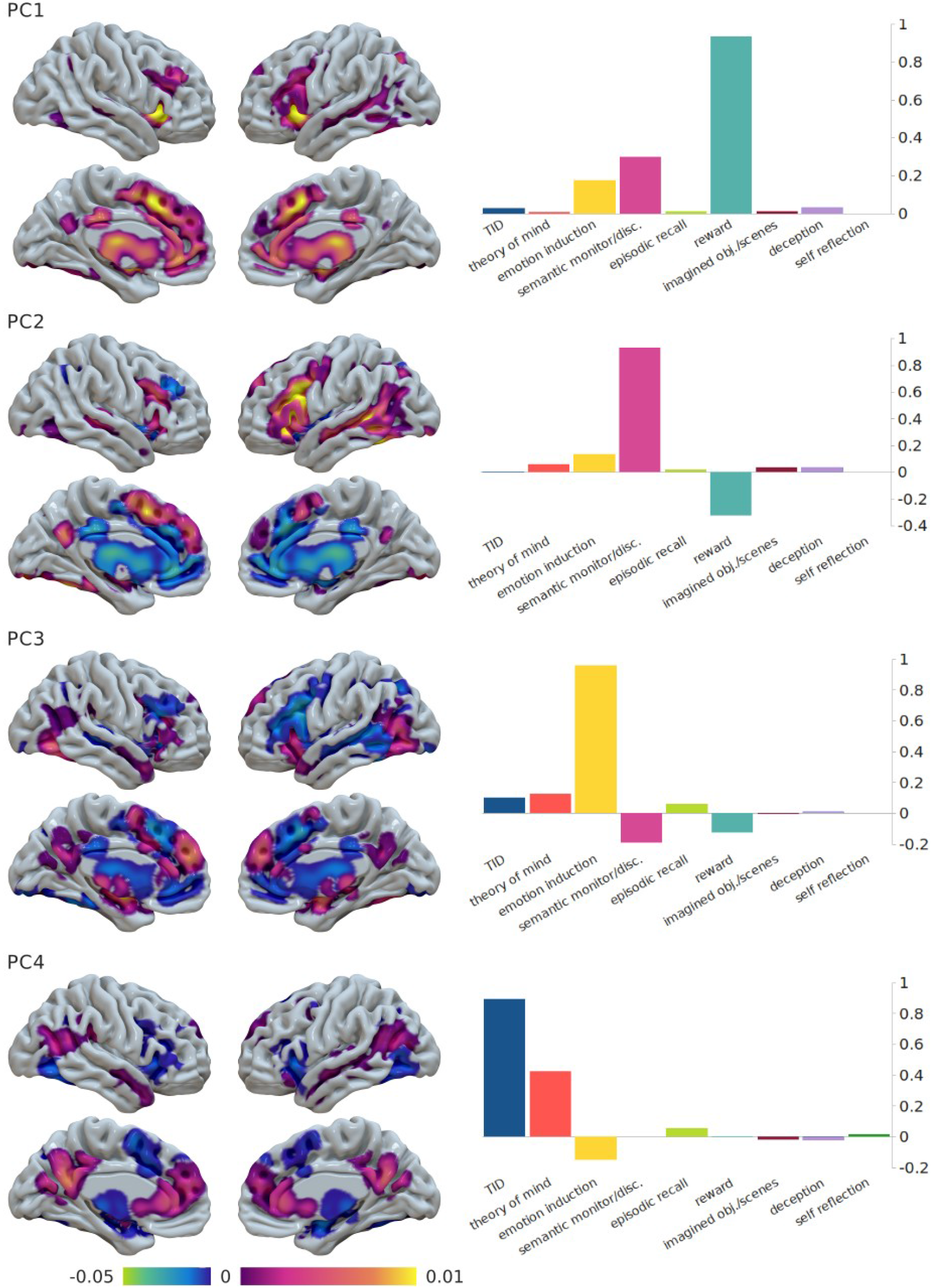
Results of the first four principal components of the Principal Component Analysis. Left: Surface mapping of the voxel-wise scores. Right: loadings of each component on each paradigm map.

The third component (PC3), which explains 10.1% of the variance and loads on the Emotion Induction map, shows a complex spatial distribution of positive and negative scores. In the mPFC, positive and negative clusters are interleaved between each other, with SMA, part of dmPFC, and associated section of the ACC showing positive values. The PCC is positive but is sided by a more anterior negative cluster. Thalamus and anterior BG are negative, but the amygdala and part of the midbrain are positive. Most lateral cortical regions, positive in PC2, are negative in PC3, except for the left AI and bilateral occipito-temporal cortex.

The first three components, taken together, explain 92.3% of the variance. The fourth one (PC4) reaches the 96.3% of cumulative explained variance. Its voxel-wise positive and negative scores show an evident similarity with the DMN and its anticorrelated network (M. D. Fox et al., 2005), and, in accordance to this, it loads on TID and ToM maps. In summary, most of our dataset variance is explained by components that load on functions not so commonly associated with the DMN (Reward and Semantic/Monitor Discrimination), with positive scores in DMN, but also SN regions. The midline core is somewhat always present, but a more conventional representation of the DMN emerges only as the fourth component. The PCA results computed excluding the Self-Reflection map were identical to the ones presented here (not shown).

### Independent Component Analysis

As the PCA indicated that four principal components provide a significant decomposition of the variance, we selected a four-component solution for the ICA as well (Fig 4). When excluding the Self-Reflection map from the data, the ICA results did not change (not shown). Since the results were similar to the ones from PCA, for an easier argumentation, we ordered the independent components so as to match the principal ones.

**Figure 4:**
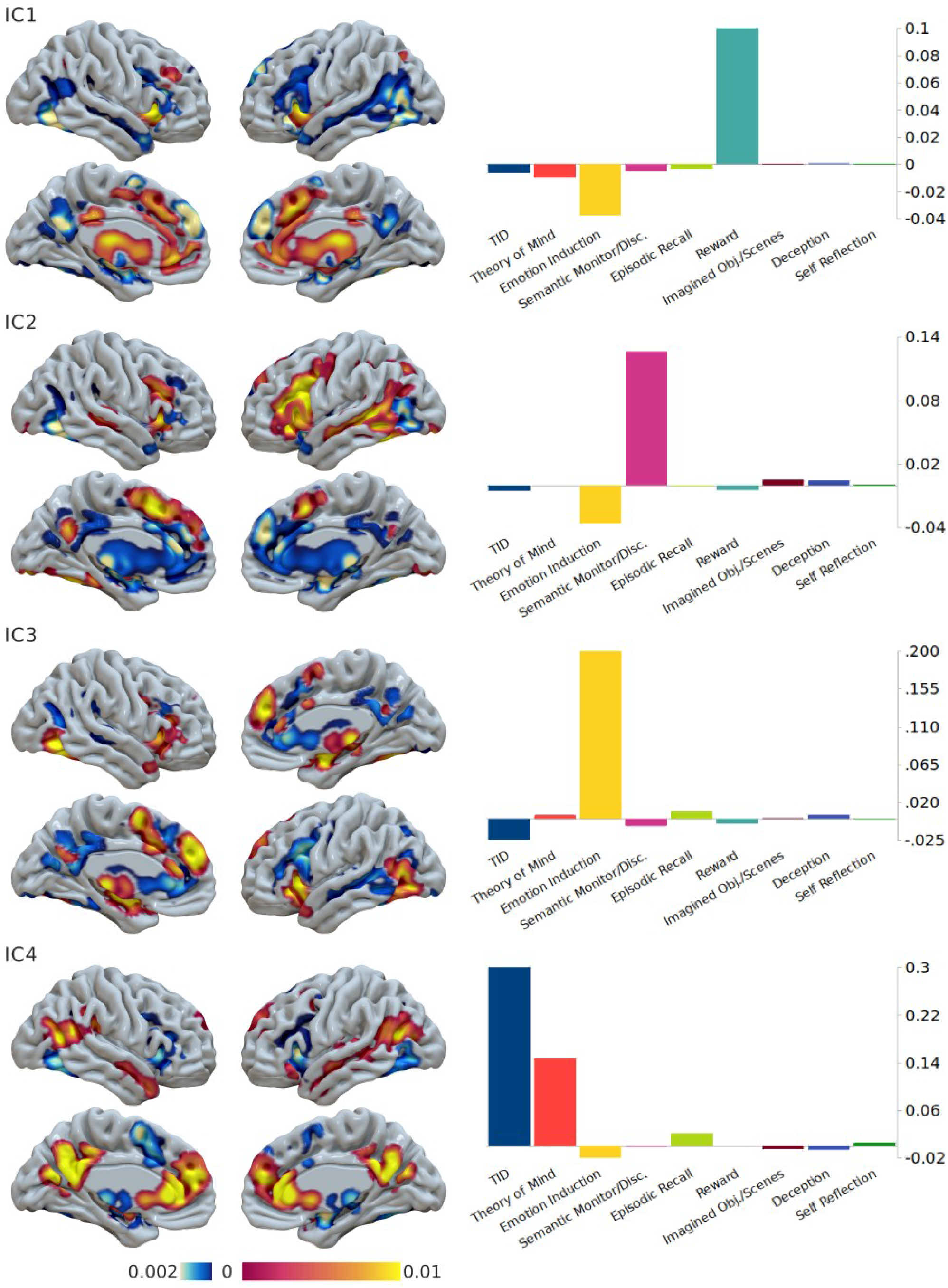
Results of the four-component solution of the Independent Component Analysis. Left: Surface mapping of the voxel-wise scores. Right: weights of the unmixing matrix of each component on each paradigm map.

IC4, just as PC1, is associated with Reward and it is positively correlated with most mPFC, anterior PCC, insula, and thalamus/BG. However, it is also anticorrelated with dmPFC, PCC proper, occipito-temporal cortices, and amygdala. This is likely because IC1, differently from PC1, has a negative load on Emotion Induction. IC2, related to Semantic Monitor/Discrimination, is similar to PC2, but in this case the dmPFC is anticorrelated, as the component loads negatively to Emotion Induction and this area is activated by such task (Fig. 1). IC3, associated with Emotion Induction, resembles PC3, but their PCC involvement is quite different. In this case, only a small PCC segment is part of the component and the rest is anticorrelated to it. IC3, like PC4, is linked to TID and ToM, it is clearly expressed in the midline core and anticorrelated with the insula, SMA, and amygdala.

Intriguingly, when the ICA maps were fed back into a Paradigm Analysis (positive voxels only), the output returned a much longer list of significant experimental tasks (Supplementary Table S1 and Figure S7), along with the paradigms heavily loaded by the component. IC4 represents an exception, as it gives only few paradigms other than those found by the three original DMN masks. This is not surprising, as IC4 is associated with TID. Since the number of voxels of the ICA masks fits within the range of the original ones (Supplementary Table 1), these results are not the product of a statistical artifact. Therefore, they suggest that IC1, IC2, and IC3 are representing modes of more extrinsic cognition compared to the canonical DMN.

It is interesting to observe that our method did not produce noise components, unlike in the case of Smith and colleagues (2009), who performed an ICA directly on the activations modeled on BrainMap foci. This could be explained as a result of ALE thresholding, which eliminated the spatial heterogeneity between experiments, as well as any unrealistic activations of some voxels obtained modeling activations as 3-D Gaussian distributions. Exploratory analyses pointed out that solutions with more than five components produced patchy maps that may be considered as noise, or also as evidence that our data have no information to be further decomposed. The five-component solution was rather similar to the one presented here, except for TID and ToM split into two different components. The ToM component involves a smaller and more dorsal mPFC cluster, and a more posterior temporo-parietal cluster than the TID one (Supplementary Fig. S5). On the contrary, solutions two and three are not particularly meaningful. They showed maps similar to the canonical DMN or its antinetwork, but their components were unable to load on more than one paradigm each (Supplementary Fig. S6). Thus, even in the absence of orthogonality constraints, the task-related modulations of the DMN defy the attempts to recapitulate their variability, providing further evidence that the internal-external DMN axis cannot be collapsed onto a singular spatial representation.

## Discussion

The present work provides compelling evidence that regions of the DMN are engaged in several tasks, which goes beyond those conventionally associated with the resting state and the mind-wandering, including also semantic reasoning and reward mechanisms. Although these tasks may be easily linked to some internal form of mentation, the nature of some of them highlights that such internal cognition seems to be crucial for external tasks. It is important to highlight that, since we tested the DMN for significant paradigms, the resulting ALE maps represent the activations of a specific set of experimental tasks, defined in an operational way. On the contrary, psychological definitions such as those implemented by the Behavioral Analysis (Lancaster et al., 2012) span across paradigms. For instance, the behavioral domain of Semantics entails Covert Word Generation, Self-Reflection, Encoding, Passive Listening, Visuospatial Attention, and other tasks. These range from the most extrinsic and active functions to forms of internal cognition that may be involved in mind-wandering. So, working at the level of paradigms allowed us to focus on the kind of activities the DMN regions are engaged in.

For instance, we found Reward paradigms to be associated with DMN regions. This is mostly due to a large amount of foci covering the whole mPFC. The latter is known to modulate reward mechanisms (Ferenczi et al., 2016), to respond to the outcome of risky decisions (Xue et al., 2009), and to activate when receiving a social reward (Martins et al., 2021). Reward mechanisms could be considered as an example of functions meant to monitor internal states, yet crucial for the implementation of behavior. As a matter of fact, reward dynamics involve the perception of somatic states and their emotional and self-related processing, which implies the activity of mPFC, ACC, SMA, along with BG, amygdala, and insula (Verdejo-García and Bechara, 2009; but see also Dunn et al., 2006). At the same time, reward functions are critical for learning, risk-taking, and behavior in general (Schultz, 2015). Furthermore, as an example of collaboration between internal and external cognition, the reward system is functionally connected to the DMN during mental simulation of the outcome of goal-directed behavior (Gerlach et al., 2014).

Deception is another paradigm class arguably standing between internal and external cognition. Deceiving someone requires social cognition, which is typically associated with the DMN internal mentation (Buckner et al., 2008). However, it might be argued that deceiving is a more external activity than ToM. While the latter only requires to represent other people’s mental states, the former also implies goal-directed programming of one’s own behavior in order to successfully deceive the other (Lisofsky et al., 2014). Moreover, specific attentional and executive functions are likely to be necessary to perform a deception paradigm (Christ et al., 2009; Farah et al., 2014). As a matter of fact, ToM appeared close to TID on the first MDS axis and Deception at the extreme opposite side.

In general, the MDS results suggested that the DMN-related experimental paradigms and their associated activation maps could be arranged along an internal-external, midline-lateral axis. This observation is consistent with the growing body of evidence pointing out that the DMN is recruited during task execution (Crittenden et al., 2015; Murphy et al., 2018; Vatansever et al., 2017), and suggests that its function may be related to some form of high-level cognition, detached from the here and now, but still crucial for goal-directed behavior (Benedek et al., 2016; Konishi et al., 2015). At the same time, our results also clearly illustrate that, when engaged in external operations, the network activations shift from the spatial representation typical of rest condition. In fact, the ALE maps associated with more extrinsic paradigms display a clear dissimilarity from the canonical representation of the DMN. Specifically, they involve peripheral nodes of the system such as AG and IFG, they often show weak or no activation at all within the midline core, and they sometimes engage SN regions considered to be anticorrelated to the DMN at rest (M. D. Fox et al., 2005)

The significance of the Paradigm Analysis on the selected tasks was likely to be driven by activations in these lateral areas. Thus, it may be argued that these maps should not be necessarily associated with the DMN, as they involve mostly its peripheral nodes as well as other external areas. However, it may also be pointed out that canonical RSNs are merely opportune simplifications of the complex, dynamic, and hierarchically multi-layered nature of FC. That is, they are heuristics helpful to reduce the connectome to a limited set of systems with taxonomic utility (Uddin et al., 2019). There are several obvious reasons explaining why canonical RSNs are a simplistic representation of brain complexity (Pessoa, 2014). For instance, they partition the gray matter into non-overlapping volumes, even if in the brain, as in most real-world networks, a node is usually connected to more than one community (Ferrarini et al., 2009; Najafi et al., 2016; Palla et al., 2005; Yeo et al., 2014). Also, FC changes from rest to task (Arbabshirani et al., 2013; Bolt et al., 2017; Cole et al., 2014; Goparaju et al., 2014; Jones et al., 2012; Krieger-Redwood et al., 2016; Mennes et al., 2013; Najafi et al., 2016; Shirer et al., 2012; Spreng et al., 2013; D. Vatansever et al., 2015; Deniz Vatansever et al., 2015) and dynamically fluctuates over time (Calhoun et al., 2014; Chang and Glover, 2010; Hutchison et al., 2013; Preti et al., 2017). The non-stationarity of FC indicates a fluid node recruitment by whole-brain connectivity modules, resulting in time-varying networks. In this regard, De Pasquale and colleagues (2012) found the DMN to be the system most connected with extra-network regions during epochs of strong internal correlation. More in general, several dynamic FC studies (Chang and Glover, 2010; Karahanoğlu and Van De Ville, 2015; Kiviniemi et al., 2011; Liu and Duyn, 2013) portrayed the DMN as a moving landscape, with a changing spatial distribution and whole-brain correlations over time. However, it might be overwhelming to deal with such degree of complexity. For this reason, it could be convenient to reduce it to a limited set of functional or structural systems (Uddin et al., 2019), upon which task-based reorganizations and functional dynamics are built (Krienen et al., 2014; Petersen and Sporns, 2015; Shine et al., 2019; Voytek and Knight, 2015).

This theoretical position presents a paradox. On one hand, it suggests that the networks we found may be more easily integrated into the current literature if seen as variations of the known template of canonical DMN, rather than as a varied set of subsisting networks. On the other hand, it highlights that none of the possible DMN variants, including the one associated here with the TID ALE, is truly representative of its dynamical activity. In other words, our view calls for a principle of epistemological simplicity, while raising the issue of ontological complexity. At the methodological level, the present work aimed to conciliate such conflicting conceptualizations by decomposing the task-related functional variance into spatial components.

Accordingly, a PCA was performed in order to define a common set of regions meant to represent the DMN across different tasks. However, the anatomical diversity of our database was clearly hard to be summarized by a single component, and the PCA was relatively inefficient as a dimensionality reduction technique. We suggest that both PC1 and PC2 represent an outward-leaning DMN, i.e., a network of areas involved in external tasks that require a certain degree of internal mentation. PC3 highlights the specific contribution of emotions to such network. Hence, large part of the task-related variance is explained in the form of proactive modes of internal cognition, in opposition to core areas active during rest and social cognition paradigms (PC4). Even under the more lenient requisite of independence, as opposed to orthogonality, the ICA seem to confirm such view. The four-component ICA almost replicated the PCA results, with the meaningful difference that the first two components were found to be anticorrelated with regions associated with Emotion Induction. Therefore, affective functions may constitute an important factor of DMN reorganization during task execution. Taken together, MDS, PCA, and ICA results indicate that the activations of DMN regions are arranged along a continuum that spans from the most internal to a more external engagement. In addition, they suggest that semantic, reward, and emotional functions may be relevant elements of such outward-leaning default-mode of cognition. Lastly, and importantly, they indicate that the modulations of the DMN activations do not converge into a representative mid-point, but rather that they somewhat gravitate around it while shifting between internal, semantic, affective or motivational modes of cognition.

An unexpected finding was that our meta-analysis was powerful enough to produce juxtapositions of components that were reminiscent of the works by Braga and colleagues (Braga et al., 2019; Braga and Buckner, 2017; DiNicola et al., 2020). As their results were originally obtained with minimally smoothed individual data, it is remarkable that something similar was achieved by our method. Another recent meta-analysis (Ngo et al., 2019) obtained a similar result, decomposing the inter-experiment DMN variability in two components. However, our methodology was able to highlight sharp contrasts between neighboring areas just analyzing the final ALE maps. This is particularly evident for PC3 and IC3, both related to Emotion Induction. In both components, the mPFC is parcellated in alternated bands of network and anti-network, with SMA and anterior dmPFC positively associated with the paradigm, and posterior dmPFC and central mPFC showing negative values. The opposite pattern was shown by IC1, related to Reward. The PCC was tightly segmented as well, particularly in IC2 and IC3, where a small section of positive voxels was surrounded by negative values. More in general, PC2, PC3, IC2, and IC3 indicated a preferential engagement of a more posterior portion of PCC in semantic monitoring and induction of emotions, with negative scores in a more anterior part. On the contrary, reward mechanisms showed the opposite pattern in IC1 and, to some extent, in PC1. Such rostro-caudal segmentation of the PCC was also observed by Leech and colleagues during the execution of an attentional task, with the caudal portion displaying less integration with the DMN and less segregation with the task-positive regions (Leech et al., 2011). To summarize, the midline core, clearly associated with TID (PC4 and IC4), appeared much more jagged in other components, presenting patterns of correlation and anticorrelation in a gradient around the corpus callosum.

### Limitations and future directions

The main limitation of this work derived from the choice of using the BrainMap database as the only source of activation foci. This was done for the sake of consistency, as the tasks significantly associated with the DMN were found using the Paradigm Analysis, which operates on the BrainMap database. Obviously, we could have integrated our data with experiments found through a systematic search on PubMed. However, a larger dataset could have possibly translated into ALE maps in disagreement with the Paradigm Analysis results, for instance without any significant activation within DMN nodes. Importantly, this would have not been necessarily due to a better representativity of the larger database, but possibly just because of a different coding of the paradigms. Having to choose between internal consistency and a larger sample size, and considering the amplitude of the functional BrainMap data archive (more than 18000 experiments in total), we preferred to conduct our whole research within the same database. This choice returned an underpowered Self-Reflection ALE. However, by removing it from the data, we obtained a similar MDS and identical PCA and ICA results. Moreover, an exploratory ALE using a liberal uncorrected threshold with p = 0.001 (not shown) revealed additional clusters in the dmPFC, mPFC, dlPFC, left insula, and IFG. A similar map would be rather consistent with our general results. Therefore, a more representative Self-Reflection map would probably be more heavily loaded by those components representing the DMN internal modes of cognition such as PC4 and IC4, rather than leading to radically different findings.

Our results indicated that the DMN is modulated by task engagement. In other words, several co-activation networks converge on the resting-state DMN nodes. Such theoretical framework would suggest a topological redefinition of the network. For instance, the Semantic Monitor/Discrimination ALE map seems to indicate that the hubness of the network has been moved from the midline core to the left IFG and middle temporal gyrus, which are peripheral nodes during rest, and to left insula and SMA, these latter part of the SN. This evidence suggests that during task execution the nodes of the DMN could update their FC and dynamically modify their topological centrality, as observed by Cole and colleagues (2013) for the frontoparietal network. However, the present work did not directly test this hypothesis. In particular, during semantic monitoring tasks, left IFG could be coupled with the middle temporal cortex, and insula with SMA, forming two relatively independent modules. Alternatively, they could be all reciprocally co-activated in a rich-club fashion. The methods used in the present research cannot disambiguate between these and other possible hypotheses. Thus, future works could be addressed toward the implementation of some methods to estimate networks of co-activations from meta-analytical data (Cauda et al., 2020; De La Vega et al., 2016; Mancuso et al., 2019; Toro et al., 2008) so as to assess the centrality of the nodes across tasks. Task-based stationary or dynamic FC could be used as well.

The present study raises a compelling question concerning the mechanism arranging the dynamical shifts from the midline core during task. An influential model proposed that the anterior insula could be responsible for coordinating the interplay between the DMN and fronto-parietal task-positive regions (Menon and Uddin, 2010; Sridharan et al., 2008). The insula was actually found by Najafi and colleagues (Najafi et al., 2016) to be connected to several modules despite a relatively low degree centrality, both during rest and emotional tasks. The AG was implicated in the same role and identified, by Kernbach and colleagues (2018), as the mediator of the interplay between different RSNs. Alternatively, the amPFC was shown to be activated during switches between stimulus-independent and stimulus-oriented thoughts (Gilbert et al., 2005), suggesting to play a role in the coordination of internal and external modes of mentation. Future studies could further investigate the issue in order to clarify which areas or mechanisms are involved in the task-based DMN re-arrangements.

### Conclusions

The present study highlights the relevance of DMN activations during the execution of tasks that are not exactly internal, nor completely external. Furthermore, it presents an anatomo-psychological gradient staring from the most internal functions, which activate the midline core, towards such relatively extrinsic mode of brain function, which involves the lateral cortices. In the light of our results, such extrinsic mode is especially related to reward, semantic, and emotional functions. Nonetheless, an important element of our results is that the spatial variability of the task-based DMN remodulations is hard to summarize. Indeed, when the brain is actively engaged in external demands, the DMN seems to be rotating on the surface of a sphere representing its multiple configurations, and none of the investigated paradigms would approach the center.

Henceforth, it could be said that the DMN considerably varies its spatial distribution during task engagement or, alternatively, that several networks of activations overlap with DMN nodes. These two views do not necessarily contradict each other and choosing one over the other is just a matter of epistemological efficacy. In both cases, they highlight that structural or resting-state scaffoldings do not suffice to explain the task-related dynamical reconfigurations of the DMN, which are arranged following a much richer functional diversity and thus showing a more spatial complexity than previously suggested.

