## Supplementary for "Default Mode Network spatial configuration varies across task domains"

### Abstract

Recent developments in network neuroscience suggest reconsidering what we thought we knew about the Default Mode Network (DMN). Although this network has always been seen as unitary and associated with the resting state, a new deconstructive line of research is pointing out that the DMN could be divided into multiple subsystems supporting different functions. By now, it is well known that the DMN is not only deactivated by tasks, but also involved in affective, mnestic, and social paradigms, among others. Nonetheless, it is starting to become clear that the array of activities in which it is involved, might also be extended to more extrinsic functions. The present meta-analytic study is meant to push this boundary a bit further. The BrainMap database was searched for all experimental paradigms activating the DMN, and their activation maps were then computed. An additional map of task-induced deactivations was also created. A Multidimensional Scaling indicated that such maps could be arranged along an anatomo-psychological gradient, which goes from midline core activations, associated with the most internal functions, to the involvement of lateral cortices in more external tasks. Further investigations suggested that such extrinsic mode is especially related to reward, semantic, and emotional functions. However, an important finding was that the variability of task-induced DMN anatomic redistribution was hard to recapitulate, as none of the maps, or any linear combination of them, could represent the whole space of its dynamical reconfiguration. Altogether, our findings suggest that the DMN may be characterized by a richer functional diversity and a more spatial complexity than previously suggested.

**Keywords:** Activation Likelihood Estimation; Multidimensional Scaling; Principal Component Analysis; Independent Component Analysis; Medial Prefrontal Cortex; Posterior Cingulate Cortex; Task Induced Deactivations; Episodic Recall; Imagination; Deception; Semantics; Reward; Theory of Mind; Emotion Induction; Self-Reflection.

*Table 1: BrainMap paradigms found to be significantly associated with the DMN masks obtained by the 7 networks atlas by Yeo et al. (2011), the Independent Component Analysis by Shirer et al. (2012), and the CAREN atlas by Doucet et al. (2019). The paradigms excluded from further analysis are written in italics.*

| **Yeo et al.** | | **Shirer et al.** | | **Doucet et al.** | |
|---|---|---|---|---|---|
| ***BrainMap paradigm*** | ***z-score*** | ***BrainMap paradigm*** | ***z-score*** | ***BrainMap paradigm*** | ***z-score*** |
| Theory of Mind | 13.696 | Theory of Mind | 7.986 | Theory of Mind | 8.359 |
| Semantic Monitor/Discrimination | 5.255 | Episodic Recall | 5.266 | Semantic Monitor/Discrimination | 6.680 |
| Episodic Recall | 5.227 | Self-Reflection | 4.395 | Episodic Recall | 5.414 |
| Emotion Induction | 4.768 | Emotion Induction | 4.262 | *Cued Explicit Recognition/Recall* | 4.369 |
| Self-Reflection | 3.872 | *Acupuncture* | 4.043 | Emotion Induction | 4.303 |
| Deception | 3.690 | Imagined Objects/Scenes | 3.954 | *Reasoning/Problem Solving* | 4.112 |
| *Passive Listening* | 3.327 | Reward | 3.458 | Reward | 3.647 |
| | | | | Deception | 3.614 |
| | | | | *Reading (Covert)* | 3.601 |
| | | | | Imagined Objects/Scenes | 3.537 |
| | | | | *Face Monitor/Discrimination* | 3.536 |

Details about the results from Sleuth searches for the 8 queries associated with each significant paradigm are presented in Table 2 and in the Prisma Flow chart in Supplementary Fig. S1. We point out that we found a limited number of experiments for the Self-Reflection condition. According to Eickhoff and colleagues (Eickhoff et al., 2016; Müller et al., 2018), at least 17 experiments should be gathered in order to perform a statistically sound ALE. Although the query returned 28 experiments, after merging them according to groups of subjects (Müller et al., 2018), they became 7 foci groupings.

To rule out possible bias induced by the inclusion of this underpowered domain, the subsequent analyses were repeated with and without the related ALE map. Since excluding this condition did not significantly influence the outcome of, we decided to keep it in the dataset.

*Table 2: Details about the result of Sleuth queries.*

| ***N experiments*** | ***N groups*** | ***N foci*** | ***N subjects*** |
|---|---|---|---|
| **Theory of Mind** | | | |
| 218 | 63 | 1663 | 1127 |
| **Semantic/Monitor Discrimination** | | | |
| 646 | 205 | 4954 | 3020 |
| **Episodic Recall** | | | |
| 123 | 39 | 1009 | 566 |
| **Emotion Induction** | | | |
| 537 | 166 | 3575 | 3234 |
| **Self-Reflection** | | | |
| 28 | 7 | 144 | 140 |
| **Deception** | | | |
| 115 | 39 | 885 | 954 |
| **Imagined Object/Scenes** | | | |
| 120 | 46 | 1097 | 660 |
| **Reward** | | | |
| 757 | 199 | 5860 | 3681 |
| **Task Induced Deactivations** | | | |
| 189 | 106 | 1665 | 1494 |

We also computed a TID meta-analysis, representing rest. The ALE results are presented in Fig. 1. The TID ALE map replicates the one by Laird and colleagues (Laird et al., 2009), with the exception of the mPFC cluster, that we found in a more dorsal position.

As for the other maps, Episodic Recall activates the left insula along with DMN areas. Imagining objects and scenes activates the left hemisphere in the precuneus, lateral prefrontal cortex (lPFC) including IFG, and supplementary motor area (SMA). Deception tasks activate the bilateral insula, other than AG and SMA, in accordance to Farah and colleagues (Farah et al., 2014). Semantic monitoring and discrimination activate the left lPFC including IFG, but also the SMA, the left lateral and medial temporal lobe, and the PCC. Reward activates the basal ganglia (BG), the thalamus, the whole mPFC, and the bilateral insula. ToM activates the temporal and temporoparietal cortices along with PCC and dmPFC, especially on the left. Inducing emotion activates the left insula and the bilateral occipito-temporal cortices, along with typical DMN nodes. The Self-Reflection map involves PCC, the right SMA, and an occipital cluster in the fusiform gyrus. The maps obtained by the Fail-safe analyses, presented in Supplementary Fig. S2, show that our ALE maps remain substantially unchanged when accounting for the file-drawer effect.

*Figure 1: Surface mapping of the 9 Activation Likelihood Estimation maps.*

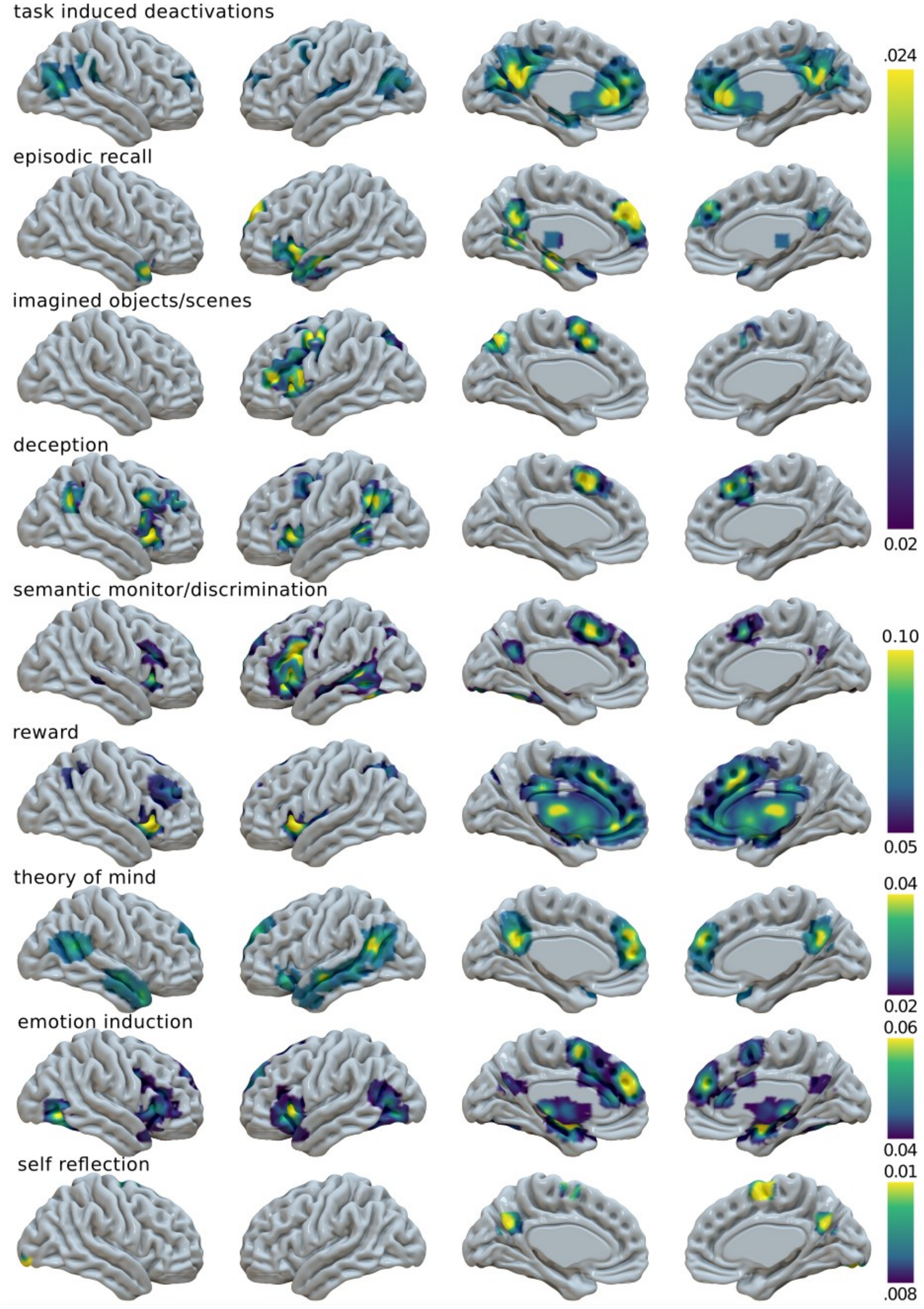

In summary, few maps matched the prototypical representation of the DMN. Some of them showed either weak activation or no activation at all in the midline core, and a strong expression of lateral areas of the network such as AG, IFG, and middle temporal gyrus. In addition, the insula and SMA/dorsal ACC, hubs of the salience network (SN), were often present. Rather than considering these as spurious findings, we see them as an indication that, when the brain is engaged by external demands, multiple networks including DMN nodes would emerge. Although relying on intrinsic brain topology, such recruitment would be not strictly constrained by it (Cole et al., 2014; Krienen et al., 2014). Thus, it might involve a flexible shift in brain hubness (Cole et al., 2013; Fransson and Thompson, 2020) and a remodulation of cooperative and competitive long-range connectivity patterns (Dixon et al., 2017; Fornito et al., 2012; Piccoli et al., 2015).

*Figure 2: Multidimensional Scaling (MDS) of the 9 Activation Likelihood Estimation (ALE) maps. Top Left: 1-correlation distance matrix of the 9 ALE maps. Top Right: MDS 2-dimensional solution. The surface maps are centered on the MDS coordinates. Bottom: MDS 3-dimensional solution, seen from different perspectives.*

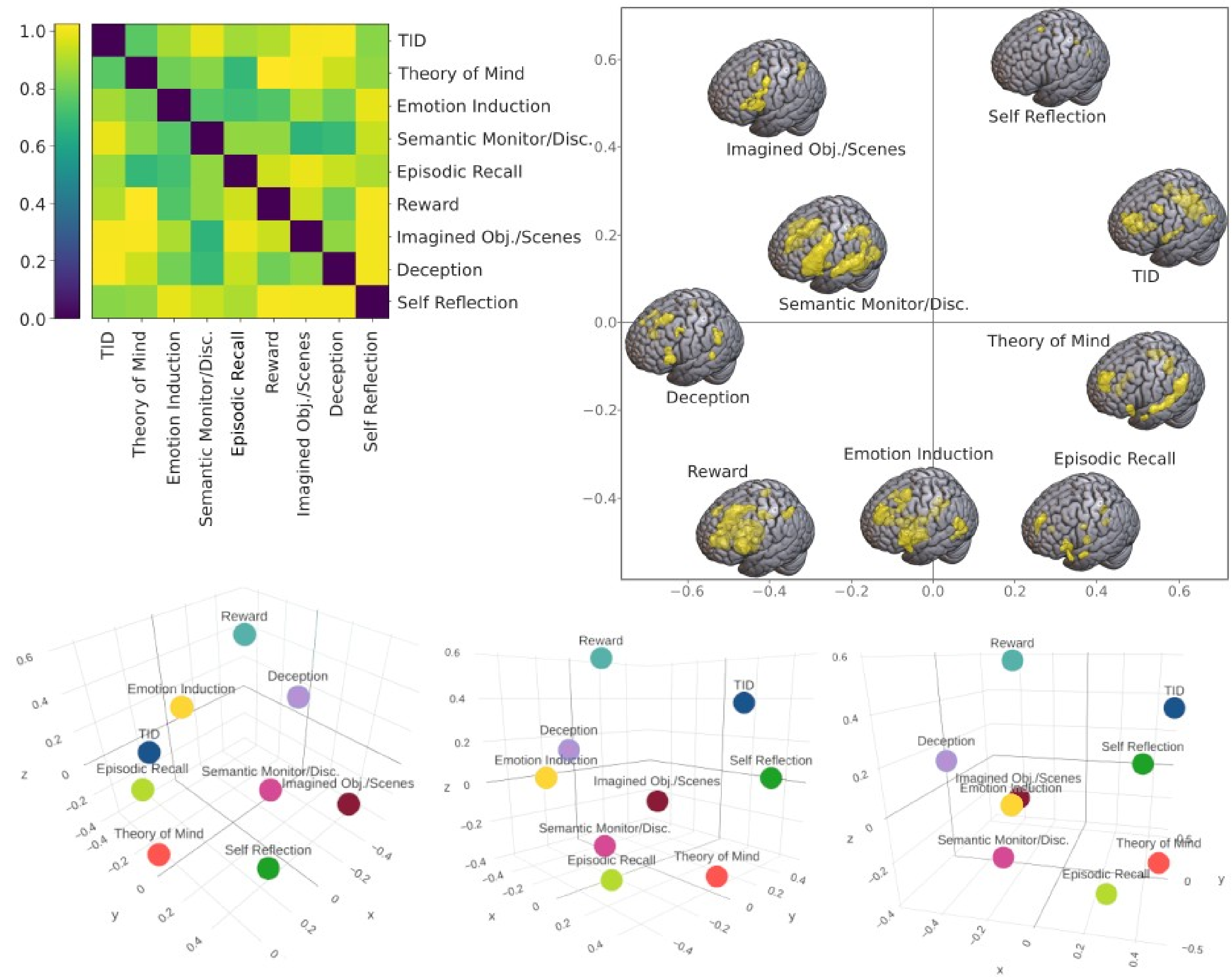

### *Principal Component Analysis*

Given the lack of common ground between different DMN expressions, we performed a PCA to summarize the inter-paradigm similarities. However, the sphericity implied by MDS raises doubts on PCA ability to reduce our data dimensionality. Even if the high significance of a Bartlett's test ($p < 0.001$) seems to suggest otherwise, the Kaiser-Meyer-Olkin index (KMO = 0.61) confirms a sufficient but mediocre relation between the maps.

*Figure 3: Results of the first four principal components of the Principal Component Analysis. Left: Surface mapping of the voxel-wise scores. Right: loadings of each component on each paradigm map.*

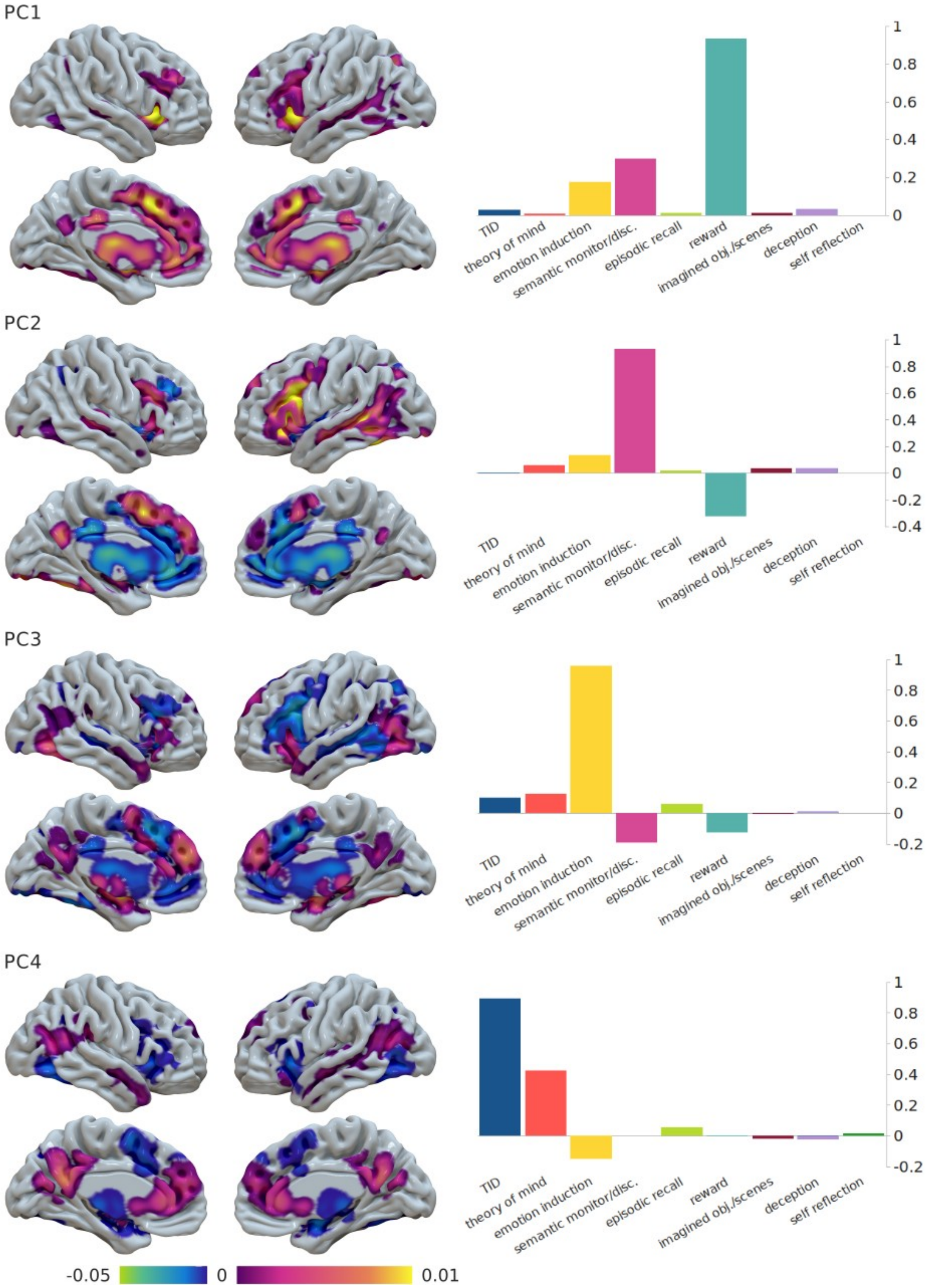

The third component (PC3), which explains 10.1% of the variance and loads on the Emotion Induction map, shows a complex spatial distribution of positive and negative scores. In the mPFC, positive and negative clusters are interleaved between each other, with SMA, part of dmPFC, and associated section of the ACC showing positive values. The PCC is positive but is sided by a more anterior negative cluster. Thalamus and anterior BG are negative, but the amygdala and part of the midbrain are positive. Most lateral cortical regions, positive in PC2, are negative in PC3, except for the left AI and bilateral occipito-temporal cortex.

The first three components, taken together, explain 92.3% of the variance. The fourth one (PC4) reaches the 96.3% of cumulative explained variance. Its voxel-wise positive and negative scores show an evident similarity with the DMN and its anticorrelated network (M. D. Fox et al., 2005), and, in accordance to this, it loads on TID and ToM maps. In summary, most of our dataset variance is explained by components that load on functions not so commonly associated with the DMN (Reward and Semantic/Monitor Discrimination), with positive scores in DMN, but also SN regions. The midline core is somewhat always present, but a more conventional representation of the DMN emerges only as the fourth component. The PCA results computed excluding the Self-Reflection map were identical to the ones presented here (not shown).

#### Independent Component Analysis

As the PCA indicated that four principal components provide a significant decomposition of the variance, we selected a four-component solution for the ICA as well (Fig 4). When excluding the Self-Reflection map from the data, the ICA results did not change (not shown). Since the results were similar to the ones from PCA, for an easier argumentation, we ordered the independent components so as to match the principal ones.

*Figure 4: Results of the four-component solution of the Independent Component Analysis. Left: Surface mapping of the voxel-wise scores. Right: weights of the unmixing matrix of each component on each paradigm map.*

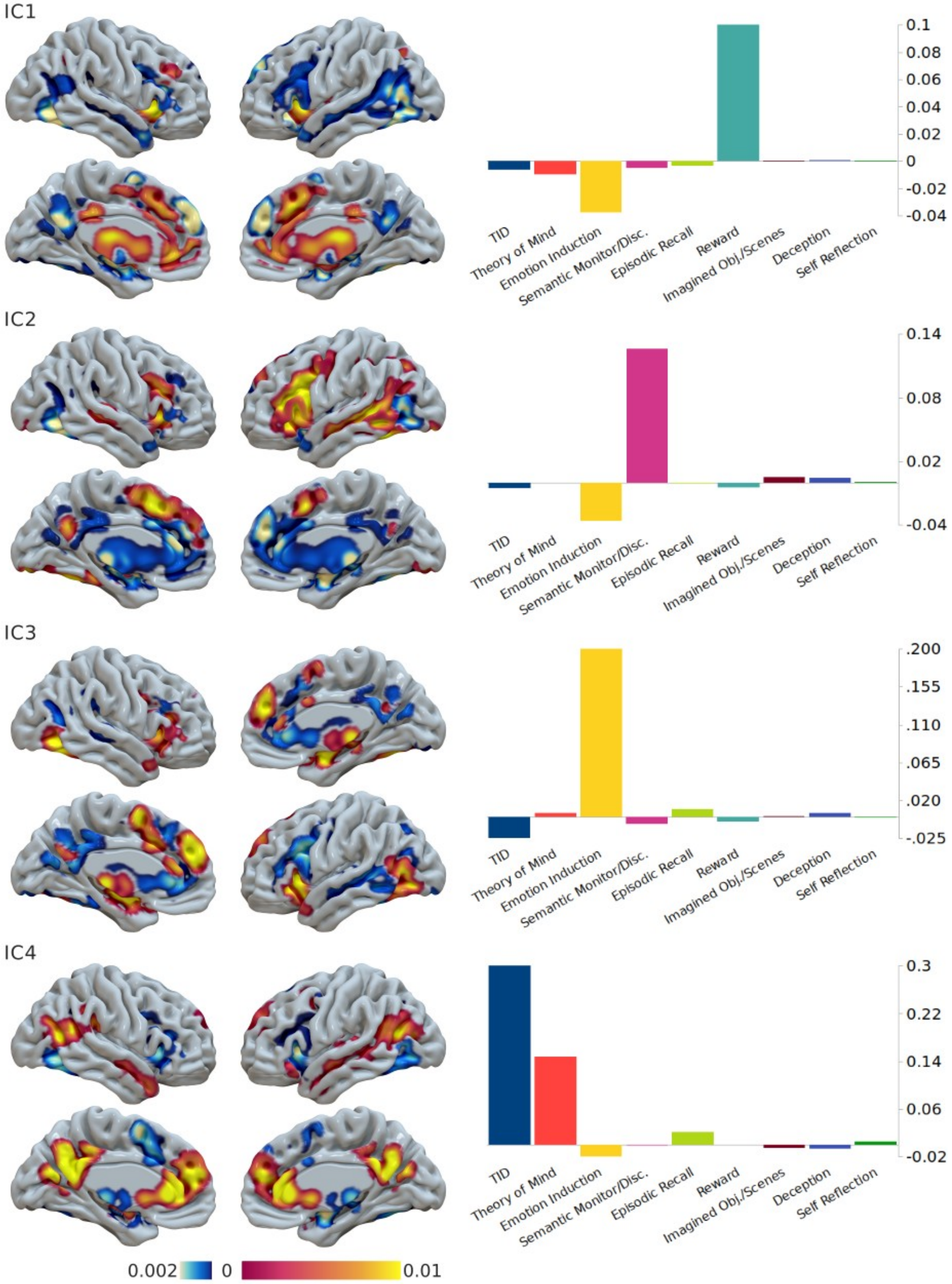

IC1
0.1
0.08
0.06
0.04
0.02
0
-0.02
-0.04
TID
Theory of Mind
Emotion Induction
Semantic Monitor/Disc.
Episodic Recall
Reward
Imagined Obj./Scenes
Deception
Self Reflection
IC2
0.14
0.08
0.02
-0.04
TID
Theory of Mind
Emotion Induction
Semantic Monitor/Disc.
Episodic Recall
Reward
Imagined Obj./Scenes
Deception
Self Reflection
IC3
.200
.155
.110
.065
.020
-.025
TID
Theory of Mind
Emotion Induction
Semantic Monitor/Disc.
Episodic Recall
Reward
Imagined Obj./Scenes
Deception
Self Reflection
IC4
0.3
0.22
0.14
0.06
-0.02
TID
Theory of Mind
Emotion Induction
Semantic Monitor/Disc.
Episodic Recall
Reward
Imagined Obj./Scenes
Deception
Self Reflection
0.002
0
0.01

Botvinik-Nezer, R., Holzmeister, F., Camerer, C.F., Dreber, A., Huber, J., Johannesson, M., Kirchler, M., Iwanir, R., Mumford, J.A., Adcock, R.A., Avesani, P., Baczkowski, B.M., Bajracharya, A., Bakst, L., Ball, S., Barilari, M., Bault, N., Beaton, D., Beitner, J., Benoit, R.G., Berkers, R.M.W.J., Bhanji, J.P., Biswal, B.B., Bobadilla-Suarez, S., Bortolini, T., Bottenhorn, K.L., Bowring, A., Braem, S., Brooks, H.R., Brudner, E.G., Calderon, C.B., Camilleri, J.A., Castrellon, J.J., Cecchetti, L., Cieslik, E.C., Cole, Z.J., Collignon, O., Cox, R.W., Cunningham, W.A., Czoschke, S., Dadi, K., Davis, C.P., Luca, A. De, Delgado, M.R., Demetriou, L., Dennison, J.B., Di, X., Dickie, E.W., Dobryakova, E., Donnat, C.L., Dukart, J., Duncan, N.W., Durnez, J., Eed, A.,

Eickhoff, S.B., Erhart, A., Fontanesi, L., Fricke, G.M., Fu, S., Galván, A., Gau, R., Genon, S., Glatard, T., Glerean, E., Goeman, J.J., Golowin, S.A.E., González-García, C., Gorgolewski, K.J., Grady, C.L., Green, M.A., Guassi Moreira, J.F., Guest, O., Hakimi, S., Hamilton, J.P., Hancock, R., Handjaras, G., Harry, B.B., Hawco, C., Herholz, P., Herman, G., Heunis, S., Hoffstaedter, F., Hogeveen, J., Holmes, S., Hu, C.-P., Huettel, S.A., Hughes, M.E., Iacovella, V., Iordan, A.D., Isager, P.M., Isik, A.I., Jahn, A., Johnson, M.R., Johnstone, T., Joseph, M.J.E., Juliano, A.C., Kable, J.W., Kassinopoulos, M., Koba, C., Kong, X.-Z., Koscik, T.R., Kucukboyaci, N.E., Kuhl, B.A., Kupek, S., Laird, A.R., Lamm, C., Langner, R., Lauharatanahirun, N., Lee, H., Lee, S., Leemans, A., Leo, A., Lesage, E., Li, F., Li, M.Y.C., Lim, P.C., Lintz, E.N., Liphardt, S.W., Losecaat Vermeer, A.B., Love, B.C., Mack, M.L., Malpica, N., Marins, T., Maumet, C., McDonald, K., McGuire, J.T., Melero, H., Méndez Leal, A.S., Meyer, B., Meyer, K.N., Mihai, G., Mitsis, G.D., Moll, J., Nielson, D.M., Nilsonne, G., Notter, M.P., Olivetti, E., Onicas, A.I., Papale, P., Patil, K.R., Peelle, J.E., Pérez, A., Pischedda, D., Poline, J.-B., Prystauka, Y., Ray, S., Reuter-Lorenz, P.A., Reynolds, R.C., Ricciardi, E., Rieck, J.R., Rodriguez-Thompson, A.M., Romyn, A., Salo, T., Samanez-Larkin, G.R., Sanz-Morales, E., Schlichting, M.L., Schultz, D.H., Shen, Q., Sheridan, M.A., Silvers, J.A., Skagerlund, K., Smith, A., Smith, D. V., Sokol-Hessner, P., Steinkamp, S.R., Tashjian, S.M., Thirion, B., Thorp, J.N., Tinghög, G., Tisdall, L., Tompson, S.H., Toro-Serey, C., Torre Tresols, J.J., Tozzi, L., Truong, V., Turella, L., van ‘t Veer, A.E., Verguts, T., Vettel, J.M., Vijayarajah, S., Vo, K., Wall, M.B., Weeda, W.D., Weis, S., White, D.J., Wisniewski, D., Xifra-Porxas, A., Yearling, E.A., Yoon, S., Yuan, R., Yuen, K.S.L., Zhang, L., Zhang, X., Zosky, J.E., Nichols, T.E., Poldrack, R.A., Schonberg, T., 2020. Variability in the analysis of a single neuroimaging dataset by many teams. Nature 582, 84–88. https://doi.org/10.1038/s41586-020-2314-9

Braga, R.M., Buckner, R.L., 2017. Parallel Interdigitated Distributed Networks within the Individual Estimated by Intrinsic Functional Connectivity. Neuron 95, 457-471.e5. https://doi.org/10.1016/j.neuron.2017.06.038

Braga, R.M., Van Dijk, K.R.A., Polimeni, J.R., Eldaief, M.C., Buckner, R.L., 2019. Parallel distributed networks resolved at high resolution reveal close juxtaposition of distinct regions. J. Neurophysiol. 121, 1513–1534. https://doi.org/10.1152/jn.00808.2018

Brett, M., Markiewicz, C.J., Hanke, M., Côté, M.-A., Cipollini, B., McCarthy, P., Jarecka, D., Cheng, C.P., Halchenko, Y.O., Cottaar, M., Larson, E., Ghosh, S., Wassermann, D., Gerhard, S., Lee, G.R., Wang, H.-T., Kastman, E., Kaczmarzyk, J., Guidotti, R., Duek, O., Daniel, J., Rokem, A., Madison, C., Moloney, B., Morency, F.C., Goncalves, M., Markello, R., Riddell, C., Burns, C., Millman, J., Gramfort, A., Leppäkangas, J., Sólon, A., van den Bosch, J.J.F., Vincent, R.D., Braun, H., Subramaniam, K., Gorgolewski, K.J., Raamana, P.R., Klug, J., Nichols, B.N., Baker, E.M., Hayashi, S., Pinsard, B., Haselgrove, C., Hymers, M., Esteban, O., Koudoro, S., Pérez-García, F., Oosterhof, N.N., Amirbekian, B., Nimmo-Smith, I., Nguyen, L., Reddigari, S., St-Jean, S., Panfilov, E., Garyfallidis, E., Varoquaux, G., Legarreta, J.H., Hahn, K.S., Hinds, O.P., Fauber, B.,

Poline, J.-B., Stutters, J., Jordan, K., Cieslak, M., Moreno, M.E., Haenel, V., Schwartz, Y., Baratz, Z., Darwin, B.C., Thirion, B., Gauthier, C., Papadopoulos Orfanos, D., Solovey, I., Gonzalez, I., Palasubramaniam, J., Lecher, J., Leinweber, K., Raktivan, K., Calábková, M., Fischer, P., Gervais, P., Gadde, S., Ballinger, T., Roos, T., Reddam, V.R., freec84, 2020. nipy/nibabel: 3.2.1. https://doi.org/10.5281/ZENODO.4295521

Buckner, R.L., Andrews-Hanna, J.R., Schacter, D.L., 2008. The brain's default network: Anatomy, function, and relevance to disease. Ann. N. Y. Acad. Sci. 1124, 1–38. https://doi.org/10.1196/annals.1440.011

Buckner, R.L., Carroll, D.C., 2007. Self-projection and the brain. Trends Cogn. Sci. 11, 49–57. https://doi.org/10.1016/j.tics.2006.11.004

Buckner, R.L., DiNicola, L.M., 2019. The brain's default network: updated anatomy, physiology and evolving insights. Nat. Rev. Neurosci. https://doi.org/10.1038/s41583-019-0212-7

Buckner, R.L., Krienen, F.M., Yeo, B.T.T., 2013. Opportunities and limitations of intrinsic functional connectivity MRI. Nat. Neurosci. 16, 832–837. https://doi.org/10.1038/nn.3423

Bzdok, D., Schilbach, L., Vogeley, K., Schneider, K., Laird, A.R., Langner, R., Eickhoff, S.B., 2012. Parsing the neural correlates of moral cognition: ALE meta-analysis on morality, theory of mind, and empathy. Brain Struct. Funct. 217, 783–796. https://doi.org/10.1007/s00429-012-0380-y

Cabeza, R., Mangels, J., Nyberg, L., Habib, R., Houle, S., McIntosh, A.R., Tulving, E., 1997. Brain regions differentially involved in remembering what and when: A PET study. Neuron 19, 863–870. https://doi.org/10.1016/S0896-6273(00)80967-8

Calhoun, V.D., Miller, R., Pearlson, G., Adali, T., 2014. The Chronnectome: Time-Varying Connectivity Networks as the Next Frontier in fMRI Data Discovery. Neuron 84, 262–274. https://doi.org/10.1016/j.neuron.2014.10.015

Cauda, F., Mancuso, L., Nani, A., Ficco, L., Premi, E., Manuello, J., Liloia, D., Gelmini, G., Duca, S., Costa, T., 2020. Hubs of long-distance co-alteration characterize brain pathology. Hum. Brain Mapp. hbm.25093. https://doi.org/10.1002/hbm.25093

Chang, C., Glover, G.H., 2010. Time-frequency dynamics of resting-state brain connectivity measured with fMRI. Neuroimage 50, 81–98. https://doi.org/10.1016/j.neuroimage.2009.12.011

Chiou, R., Humphreys, G.F., Lambon Ralph, M.A., 2020. Bipartite functional fractionation within the default network supports disparate forms of internally oriented cognition. Cereb. Cortex 30, 5484–5501. https://doi.org/10.1093/cercor/bhaa130

Christ, S.E., Van Essen, D.C., Watson, J.M., Brubaker, L.E., McDermott, K.B., 2009. The contributions of prefrontal cortex and executive control to deception: Evidence from activation likelihood estimate meta-analyses. Cereb. Cortex 19, 1557–1566. https://doi.org/10.1093/cercor/bhn189

Christoff, K., Irving, Z.C., Fox, K.C.R., Spreng, R.N., Andrews-Hanna, J.R., 2016. Mind-wandering as spontaneous thought: A dynamic framework. Nat. Rev. Neurosci. 17, 718–731. https://doi.org/10.1038/nrn.2016.113

Cocchi, L., Zalesky, A., Fornito, A., Mattingley, J.B., 2013. Dynamic cooperation and competition between brain systems during cognitive control. Trends Cogn. Sci. 17, 493–501. https://doi.org/10.1016/j.tics.2013.08.006

Cole, M.W., Bassett, D.S., Power, J.D., Braver, T.S., Petersen, S.E., 2014. Intrinsic and task-evoked network architectures of the human brain. Neuron 83, 238–251. https://doi.org/10.1016/j.neuron.2014.05.014

Cole, M.W., Reynolds, J.R., Power, J.D., Repovs, G., Anticevic, A., Braver, T.S., 2013. Multi-task connectivity reveals flexible hubs for adaptive task control. Nat. Neurosci. 16, 1348–1355. https://doi.org/10.1038/nn.3470

Crittenden, B.M., Mitchell, D.J., Duncan, J., 2015. Recruitment of the default mode network during a demanding act of executive control. Elife 2015, 1–12. https://doi.org/10.7554/eLife.06481

D'Argembeau, A., Stawarczyk, D., Majerus, S., Collette, F., Van Der Linden, M., Feyers, D., Maquet, P., Salmon, E., 2010. The neural basis of personal goal processing when envisioning future events. J. Cogn. Neurosci. 22, 1701–1713. https://doi.org/10.1162/jocn.2009.21314

Damoiseaux, J.S., Rombouts, S.A.R.B., Barkhof, F., Scheltens, P., Stam, C.J., Smith, S.M., Beckmann, C.F., 2006. Consistent resting-state networks across healthy subjects. Proc. Natl. Acad. Sci. 103, 13848–13853. https://doi.org/10.1073/pnas.0601417103

De La Vega, A., Chang, L.J., Banich, M.T., Wager, T.D., Yarkoni, T., 2016. Large-scale meta-analysis of human medial frontal cortex reveals tripartite functional organization. J. Neurosci. 36, 6553–6562. https://doi.org/10.1523/JNEUROSCI.4402-15.2016

De Luca, M., Beckmann, C.F., De Stefano, N., Matthews, P.M., Smith, S.M., 2006. fMRI resting state networks define distinct modes of long-distance interactions in the human brain. Neuroimage 29, 1359–1367. https://doi.org/10.1016/j.neuroimage.2005.08.035

de Pasquale, F., Della Penna, S., Snyder, A.Z., Marzetti, L., Pizzella, V., Romani, G.L., Corbetta, M., 2012. A Cortical Core for Dynamic Integration of Functional Networks in the Resting Human Brain. Neuron 74, 753–764. https://doi.org/10.1016/j.neuron.2012.03.031

Delgado, M.R., Beer, J.S., Fellows, L.K., Huettel, S.A., Platt, M.L., Quirk, G.J., Schiller, D., 2016. Viewpoints: Dialogues on the functional role of the ventromedial prefrontal cortex. Nat. Neurosci. 19, 1545–1552. https://doi.org/10.1038/nn.4438

Denkova, E., Nomi, J.S., Uddin, L.Q., Jha, A.P., 2019. Dynamic brain network configurations during rest and an attention task with frequent occurrence of mind wandering. Hum. Brain Mapp. 40, 4564–4576. https://doi.org/10.1002/hbm.24721

Denny, B.T., Kober, H., Wager, T.D., Ochsner, K.N., 2012. A meta-analysis of functional neuroimaging studies of self- and other judgments reveals a spatial gradient for mentalizing in medial prefrontal cortex. J. Cogn. Neurosci. 24, 1742–1752. https://doi.org/10.1162/jocn_a_00233

DiNicola, L.M., Braga, R.M., Buckner, R.L., 2020. Parallel distributed networks dissociate episodic and social functions within the individual. J. Neurophysiol. 123, 1144–1179. https://doi.org/10.1152/jn.00529.2019

Dixon, M.L., Andrews-Hanna, J.R., Spreng, R.N., Irving, Z.C., Mills, C., Girn, M., Christoff, K., 2017. Interactions between the default network and dorsal attention network vary across default subsystems, time, and cognitive states. Neuroimage 147, 632–649. https://doi.org/10.1016/j.neuroimage.2016.12.073

Dixon, M.L., Fox, K.C.R., Christoff, K., 2014. A framework for understanding the relationship between externally and internally directed cognition. Neuropsychologia 62, 321–330. https://doi.org/10.1016/j.neuropsychologia.2014.05.024

Doucet, G.E., Lee, W.H., Frangou, S., 2019. Evaluation of the spatial variability in the major resting-state networks across human brain functional atlases. Hum. Brain Mapp. hbm.24722. https://doi.org/10.1002/hbm.24722

Dunn, B.D., Dalgleish, T., Lawrence, A.D., 2006. The somatic marker hypothesis: A critical evaluation. Neurosci. Biobehav. Rev. 30, 239–271. https://doi.org/10.1016/j.neubiorev.2005.07.001

Eichele, T., Debener, S., Calhoun, V.D., Specht, K., Engel, A.K., Hugdahl, K., Von Cramon, D.Y., Ullsperger, M., 2008. Prediction of human errors by maladaptive changes in event-related brain networks. Proc. Natl. Acad. Sci. U. S. A. 105, 6173–6178. https://doi.org/10.1073/pnas.0708965105

Eickhoff, S., Laird, A., Grefkes, C., Wang, L.E., Zilles, K., Fox, P.T., 2009. Coordinate-based ALE meta-analysis of neuroimaging data: a random-effects approach based on empirical estimates of spatial uncertainty. Hum. Brain Mapp. 30, 2907–2926. https://doi.org/10.1002/hbm.20718

Eickhoff, S.B., Bzdok, D., Laird, A.R., Kurth, F., Fox, P.T., 2012. Activation likelihood estimation meta-analysis revisited. Neuroimage 59, 2349–2361. https://doi.org/10.1016/j.neuroimage.2011.09.017

Eickhoff, S.B., Nichols, T.E., Laird, A.R., Hoffstaedter, F., Amunts, K., Fox, P.T., Bzdok, D., Eickhoff, C.R., 2016. Behavior, sensitivity, and power of activation likelihood estimation characterized by massive empirical simulation. Neuroimage 137, 70–85. https://doi.org/10.1016/j.neuroimage.2016.04.072

Eickhoff, S.B., Thirion, B., Varoquaux, G., Bzdok, D., 2015. Connectivity-based parcellation: Critique and implications. Hum. Brain Mapp. 36, 4771–4792. https://doi.org/10.1002/hbm.22933

Ellamil, M., Dobson, C., Beeman, M., Christoff, K., 2012. Evaluative and generative modes of thought during the creative process. Neuroimage 59, 1783–1794. https://doi.org/10.1016/j.neuroimage.2011.08.008

Elton, A., Gao, W., 2015. Task-positive Functional Connectivity of the Default Mode Network Transcends Task Domain. J. Cogn. Neurosci. 27, 2369–2381. https://doi.org/10.1162/jocn_a_00859

Engen, H.G., Kanske, P., Singer, T., 2017. The neural component-process architecture of endogenously generated emotion. Soc. Cogn. Affect. Neurosci. 12, 197–211. https://doi.org/10.1093/scan/nsw108

Evans, M., Krieger-Redwood, K., Gonzalez Alam, T.R., Smallwood, J., Jefferies, E., 2020. Controlled semantic summation correlates with intrinsic connectivity between default mode and control networks. Cortex 129, 356–375. https://doi.org/10.1016/j.cortex.2020.04.032

Farah, M.J., Hutchinson, J.B., Phelps, E.A., Wagner, A.D., 2014. Functional MRI-based lie detection: Scientific and societal challenges. Nat. Rev. Neurosci. 15, 123–131. https://doi.org/10.1038/nrn3665

Ferenczi, E.A., Zalocusky, K.A., Liston, C., Grosenick, L., Warden, M.R., Amatya, D., Katovich, K., Mehta, H., Patenaude, B., Ramakrishnan, C., Kalanithi, P., Etkin, A., Knutson, B., Glover, G.H., Deisseroth, K., 2016. Prefrontal cortical regulation of brainwide circuit dynamics and reward-related behavior. Science (80-. ). https://doi.org/10.1126/science.aac9698

Ferrarini, L., Veer, I.M., Baerends, E., van Tol, M.-J., Renken, R.J., van der Wee, N.J.A., Veltman, D.J., Aleman, A., Zitman, F.G., Penninx, B.W.J.H., van Buchem, M.A., Reiber, J.H.C., Rombouts, S.A.R.B., Milles, J., 2009. Hierarchical functional modularity in the resting-state human brain. Hum. Brain Mapp. 30, 2220–2231. https://doi.org/10.1002/hbm.20663

Fingelkurts, Andrew A., Fingelkurts, Alexander A., Kallio-Tamminen, T., 2020. Selfhood triumvirate: From phenomenology to brain activity and back again. Conscious. Cogn. 86, 103031. https://doi.org/10.1016/j.concog.2020.103031

Fornito, A., Harrison, B.J., Zalesky, A., Simons, J.S., 2012. Competitive and cooperative dynamics of large-scale brain functional networks supporting recollection. Proc. Natl. Acad. Sci. U. S. A. 109, 12788–12793. https://doi.org/10.1073/pnas.1204185109

Fossati, P., Hevenor, S.J., Graham, S.J., Grady, C., Keightley, M.L., Craik, F., Mayberg, H., 2003. In search of the emotional self: An fMRI study using positive and negative emotional words. Am. J. Psychiatry 160, 1938–1945. https://doi.org/10.1176/appi.ajp.160.11.1938

Foster, B.L., Dastjerdi, M., Parvizi, J., 2012. Neural populations in human posteromedial cortex display opposing responses during memory and numerical processing. Proc. Natl. Acad. Sci. U. S. A. 109, 15514–15519. https://doi.org/10.1073/pnas.1206580109

Fox, K.C.R., Spreng, R.N., Ellamil, M., Andrews-Hanna, J.R., Christoff, K., 2015. The wandering brain: Meta-analysis of functional neuroimaging studies of mind-wandering and related spontaneous thought processes. Neuroimage 111, 611–621. https://doi.org/10.1016/j.neuroimage.2015.02.039

Fox, M.D., Snyder, A.Z., Vincent, J.L., Corbetta, M., Van Essen, D.C., Raichle, M.E., 2005. The human brain is intrinsically organized into dynamic, anticorrelated functional networks. Proc. Natl. Acad. Sci. 102, 9673–9678. https://doi.org/10.1073/pnas.0504136102

Fox, P.T., Laird, A.R., Fox, S.P., Fox, P.M., Uecker, A.M., Crank, M., Koenig, S.F., Lancaster, J.L., 2005. BrainMap taxonomy of experimental design: Description and evaluation. Hum. Brain Mapp. 25, 185–198. https://doi.org/10.1002/hbm.20141

Fox, P.T., Lancaster, J.L., 2002. Mapping context and content: The BrainMap model. Nat. Rev. Neurosci. 3, 319–321. https://doi.org/10.1038/nrn789

Fransson, P., 2005. Spontaneous low-frequency BOLD signal fluctuations: An fMRI investigation of the resting-state default mode of brain function hypothesis. Hum. Brain Mapp. 26, 15–29. https://doi.org/10.1002/hbm.20113

Fransson, P., Thompson, W.H., 2020. Temporal flow of hubs and connectivity in the human brain. Neuroimage 223, 117348. https://doi.org/10.1016/j.neuroimage.2020.117348

Gerlach, K.D., Spreng, R.N., Gilmore, A.W., Schacter, D.L., 2011. Solving future problems: Default network and executive activity associated with goal-directed mental simulations. Neuroimage 55, 1816–1824. https://doi.org/10.1016/j.neuroimage.2011.01.030

Gerlach, K.D., Spreng, R.N., Madore, K.P., Schacter, D.L., 2014. Future planning: default network activity couples with frontoparietal control network and reward-processing regions during process and outcome simulations. Soc. Cogn. Affect. Neurosci. 9, 1942–1951. https://doi.org/10.1093/scan/nsu001

Giambra, L.M., 1989. Task-unrelated thought frequency as a function of age: A laboratory study. Psychol. Aging 4, 136–143. https://doi.org/10.1037/0882-7974.4.2.136

Gilbert, S.J., Dumontheil, I., Simons, J.S., Frith, C.D., Burgess, P.W., 2007. Comment on "Wandering Minds: The Default Network and Stimulus-Independent Thought." Science (80-. ). 317, 43b-43b. https://doi.org/10.1126/science.1140801

Gilbert, S.J., Frith, C.D., Burgess, P.W., 2005. Involvement of rostral prefrontal cortex in selection between stimulus-oriented and stimulus-independent thought. Eur. J. Neurosci. 21, 1423–1431. https://doi.org/10.1111/j.1460-9568.2005.03981.x

Gilbert, S.J., Simons, J.S., Frith, C.D., Burgess, P.W., 2006. Performance-related activity in medial rostral prefrontal cortex (area 10) during low-demand tasks. J. Exp. Psychol. Hum. Percept. Perform. 32, 45–58. https://doi.org/10.1037/0096-1523.32.1.45

Golland, Y., Bentin, S., Gelbard, H., Benjamini, Y., Heller, R., Nir, Y., Hasson, U., Malach, R., 2007. Extrinsic and intrinsic systems in the posterior cortex of the human brain revealed during natural sensory stimulation. Cereb. Cortex 17, 766–777. https://doi.org/10.1093/cercor/bhk030

Goparaju, B., Rana, K.D., Calabro, F.J., Vaina, L.M., 2014. A computational study of whole-brain connectivity in resting state and task fMRI. Med. Sci. Monit. 20, 1024–1042. https://doi.org/10.12659/MSM.891142

Gordon, E.M., Laumann, T.O., Marek, S., Raut, R. V, Gratton, C., Newbold, D.J., Greene, D.J., Coalson, R.S., Snyder, A.Z., Schlaggar, B.L., Petersen, S.E., Dosenbach, N.U.F., Nelson, S.M., 2020. Default-mode network streams for coupling to language and control systems. Proc. Natl. Acad. Sci. U. S. A. 1–12. https://doi.org/10.1073/pnas.2005238117

Greene, J.D., Sommerville, R.B., Nystrom, L.E., Darley, J.M., Cohen, J.D., 2001. An fMRI investigation of emotional engagement in moral judgment. Science (80-. ). 293, 2105–2108. https://doi.org/10.1126/science.1062872

Greicius, M.D., Krasnow, B., Reiss, A.L., Menon, V., 2003. Functional connectivity in the resting brain: A network analysis of the default mode hypothesis. Proc. Natl. Acad. Sci. 100, 253–258. https://doi.org/10.1073/pnas.0135058100

Greicius, M.D., Menon, V., 2004. Default-Mode Activity during a Passive Sensory Task: Uncoupled from Deactivation but Impacting Activation. J. Cogn. Neurosci. 16, 1484–1492. https://doi.org/10.1162/0898929042568532

Gusnard, D. a, Raichle, M.E., 2001. Searching for a baseline: Functional imaging and the resting human brain. Nat. Rev. Neurosci. 2, 685–694. https://doi.org/10.1038/35094500

Gusnard, D.A., Akbudak, E., Shulman, G.L., Raichle, M.E., 2001. Medial prefrontal cortex and self-referential mental activity: relation to a default mode of brain function. Pnas 98, 4259–4264. https://doi.org/10.1073/pnas.071043098

Hampson, M., Driesen, N.R., Skudlarski, P., Gore, J.C., Constable, R.T., 2006. Brain connectivity related to working memory performance. J. Neurosci. 26, 13338–13343. https://doi.org/10.1523/JNEUROSCI.3408-06.2006

Harris, C.R., Millman, K.J., van der Walt, S.J., Gommers, R., Virtanen, P., Cournapeau, D., Wieser, E., Taylor, J., Berg, S., Smith, N.J., Kern, R., Picus, M., Hoyer, S., van Kerkwijk, M.H., Brett, M., Haldane, A., del Río, J.F., Wiebe, M., Peterson, P., Gérard-Marchant, P., Sheppard, K., Reddy, T., Weckesser, W., Abbasi, H., Gohlke, C., Oliphant, T.E., 2020. Array programming with NumPy. Nature 585, 357–362. https://doi.org/10.1038/s41586-020-2649-2

Harrison, B.J., Pujol, J., López-Solà, M., Hernández-Ribas, R., Deus, J., Ortiz, H., Soriano-Mas, C., Yücel, M., Pantelis, C., Cardoner, N., 2008. Consistency and functional specialization in the default mode brain network. Proc. Natl. Acad. Sci. U. S. A. 105, 9781–9786. https://doi.org/10.1073/pnas.0711791105

Hassabis, D., Kumaran, D., Maguire, E.A., 2007. Using imagination to understand the neural basis of episodic memory. J. Neurosci. 27, 14365–14374. https://doi.org/10.1523/JNEUROSCI.4549-07.2007

Hassabis, D., Maguire, E.A., 2007. Deconstructing episodic memory with construction. Trends Cogn. Sci. 11, 299–306. https://doi.org/10.1016/j.tics.2007.05.001

Hiser, J., Koenigs, M., 2018. The Multifaceted Role of the Ventromedial Prefrontal Cortex in Emotion, Decision Making, Social Cognition, and Psychopathology. Biol. Psychiatry 83, 638–647. https://doi.org/10.1016/j.biopsych.2017.10.030

Huo, T., Li, Y., Zhuang, K., Song, L., Wang, X., Ren, Z., Liu, Q., Yang, W., Qiu, J., 2020. Industriousness Moderates the Link Between Default Mode Network Subsystem and Creativity. Neuroscience. https://doi.org/10.1016/j.neuroscience.2019.11.049

Hutchison, R.M., Womelsdorf, T., Allen, E.A., Bandettini, P.A., Calhoun, V.D., Corbetta, M., Della Penna, S., Duyn, J.H., Glover, G.H., Gonzalez-Castillo, J., Handwerker, D.A., Keilholz, S., Kiviniemi, V., Leopold, D.A., de Pasquale, F., Sporns, O., Walter, M., Chang, C., 2013. Dynamic functional connectivity: Promise, issues, and interpretations. Neuroimage 80, 360–378. https://doi.org/10.1016/j.neuroimage.2013.05.079

Hyvarinen, A., 1999. Fast and robust fixed-point algorithms for independent component analysis. IEEE Trans. Neural Networks 10, 626–634. https://doi.org/10.1109/72.761722

Hyvärinen, A., Oja, E., 1997. A Fast Fixed-Point Algorithm for Independent Component Analysis. Neural Comput. 9, 1483–1492. https://doi.org/10.1162/neco.1997.9.7.1483

Jenkinson, M., Bannister, P., Brady, M., Smith, S., 2002. Improved Optimization for the Robust and Accurate Linear Registration and Motion Correction of Brain Images. Neuroimage 17, 825–841. https://doi.org/https://doi.org/10.1006/nimg.2002.1132

Jones, D.T., Vemuri, P., Murphy, M.C., Gunter, J.L., Senjem, M.L., Machulda, M.M., Przybelski, S.A., Gregg, B.E., Kantarci, K., Knopman, D.S., Boeve, B.F., Petersen, R.C., Jack, C.R., 2012. Non-stationarity in the “resting brain’s” modular architecture. PLoS One 7. https://doi.org/10.1371/journal.pone.0039731

Jung, R.E., Mead, B.S., Carrasco, J., Flores, R.A., 2013. The structure of creative cognition in the human brain. Front. Hum. Neurosci. https://doi.org/10.3389/fnhum.2013.00330

Kam, J.W.Y., Dao, E., Farley, J., Fitzpatrick, K., Smallwood, J., Schooler, J.W., Handy, T.C., 2011. Slow Fluctuations in Attentional Control of Sensory Cortex. J. Cogn. Neurosci. 23, 460–470. https://doi.org/10.1162/jocn.2010.21443

Kam, J.W.Y., Handy, T.C., 2013. The neurocognitive consequences of the wandering mind: a mechanistic account of sensory-motor decoupling. Front. Psychol. 4, 1–13. https://doi.org/10.3389/fpsyg.2013.00725

Karahanoğlu, F.I., Van De Ville, D., 2015. Transient brain activity disentangles fMRI resting-state dynamics in terms of spatially and temporally overlapping networks. Nat. Commun. 6, 7751. https://doi.org/10.1038/ncomms8751

Kernbach, J.M., Thomas Yeo, B.T., Smallwood, J., Margulies, D.S., De Schotten, M.T., Walter, H., Sabuncu, M.R., Holmes, A.J., Gramfort, A., Varoquaux, G., Thirion, B., Bzdok, D., 2018. Subspecialization within default mode nodes characterized in 10,000 UK Biobank participants. Proc. Natl. Acad. Sci. U. S. A. 115, 12295–12300. https://doi.org/10.1073/pnas.1804876115

Kim, H., 2016. Default network activation during episodic and semantic memory retrieval: A selective meta-analytic comparison. Neuropsychologia 80, 35–46. https://doi.org/10.1016/j.neuropsychologia.2015.11.006

Kiviniemi, V., Vire, T., Remes, J., Elseoud, A.A., Starck, T., Tervonen, O., Nikkinen, J., 2011. A Sliding Time-Window ICA Reveals Spatial Variability of the Default Mode Network in Time. Brain Connect. 1, 339–347. https://doi.org/10.1089/brain.2011.0036

Knyazev, G.G., Savostyanov, A.N., Bocharov, A. V., Levin, E.A., Rudych, P.D., 2020. Intrinsic Connectivity Networks in the Self- and Other-Referential Processing. Front. Hum. Neurosci. 14. https://doi.org/10.3389/fnhum.2020.579703

Konishi, M., McLaren, D.G., Engen, H., Smallwood, J., 2015. Shaped by the past: The default mode network supports cognition that is independent of immediate perceptual input. PLoS One 10, 1–18. https://doi.org/10.1371/journal.pone.0132209

Koshino, H., Minamoto, T., Ikeda, T., Osaka, M., Otsuka, Y., Osaka, N., 2011. Anterior medial prefrontal cortex exhibits activation during task preparation but deactivation during task execution. PLoS One 6, 54–58. https://doi.org/10.1371/journal.pone.0022909

Koshino, H., Minamoto, T., Yaoi, K., Osaka, M., Osaka, N., 2014. Coactivation of the default mode network regions and working memory network regions during task preparation. Sci. Rep. 4, 34–39. https://doi.org/10.1038/srep05954

Kounios, J., Frymiare, J.L., Bowden, E.M., Fleck, J.I., Subramaniam, K., Parrish, T.B., Jung-Beeman, M., 2006. The prepared mind: Neural activity prior to problem presentation predicts subsequent solution by sudden insight. Psychol. Sci. https://doi.org/10.1111/j.1467-9280.2006.01798.x

Krieger-Redwood, K., Jefferies, E., Karapanagiotidis, T., Seymour, R., Nunes, A., Ang, J.W.A., Majernikova, V., Mollo, G., Smallwood, J., 2016. Down but not out in posterior cingulate cortex: Deactivation yet functional coupling with prefrontal cortex during demanding semantic cognition. Neuroimage 141, 366–377. https://doi.org/10.1016/j.neuroimage.2016.07.060

Kriegeskorte, N., Mur, M., Bandettini, P., 2008. Representational similarity analysis - connecting the branches of systems neuroscience. Front. Syst. Neurosci. 2, 1–28. https://doi.org/10.3389/neuro.06.004.2008

Krienen, F.M., Thomas Yeo, B.T., Buckner, R.L., Yeo, B.T.T., Buckner, R.L., 2014. Reconfigurable task-dependent functional coupling modes cluster around a core functional architecture. Philos. Trans. R. Soc. B Biol. Sci. 369, 20130526–20130526. https://doi.org/10.1098/rstb.2013.0526

Laird, A.R., Eickhoff, S.B., Li, K., Robin, D.A., Glahn, D.C., Fox, P.T., 2009. Investigating the functional heterogeneity of the DMN using coordinate base metaanalytic modeling. J. Neurosci. 29, 14496–14505. https://doi.org/10.1523/JNEUROSCI.4004-09.2009.Investigating

Laird, A.R., Fox, P.M., Eickhoff, S.B., Turner, J.A., Ray, K.L., McKay, D.R., Glahn, D.C., Beckmann, C.F., Smith, S.M., Fox, P.T., 2011. Behavioral Interpretations of Intrinsic Connectivity Networks. J. Cogn. Neurosci. 23, 4022–4037. https://doi.org/10.1162/jocn_a_00077

Laird, A.R., Lancaster, J.L., Fox, P.T., 2005. BrainMap: The Social Evolution of a Human Brain Mapping Database. Neuroinformatics 3, 065–078. https://doi.org/10.1385/NI:3:1:065

Lancaster, J.L., Cykowski, M.D., McKay, D.R., Kochunov, P. V., Fox, P.T., Rogers, W., Toga, A.W., Zilles, K., Amunts, K., Mazziotta, J., 2010. Anatomical global spatial normalization. Neuroinformatics 8, 171–182. https://doi.org/10.1007/s12021-010-9074-x

Lancaster, J.L., Laird, A.R., Eickhoff, S.B., Martinez, M.J., Fox, P.M., Fox, P.T., 2012. Automated regional behavioral analysis for human brain images. Front. Neuroinform. 6, 1–12. https://doi.org/10.3389/fninf.2012.00023

Lancaster, J.L., McKay, D.R., Cykowski, M.D., Martinez, M.J., Tan, X., Valaparla, S., Zhang, Y., Fox, P.T., 2011. Automated Analysis of Fundamental Features of Brain Structures. Neuroinformatics 9, 371–380. https://doi.org/10.1007/s12021-011-9108-z

Lanzoni, L., Ravasio, D., Thompson, H., Vatansever, D., Margulies, D., Smallwood, J., Jefferies, E., 2020. The role of default mode network in semantic cue integration. Neuroimage 219, 117019. https://doi.org/10.1016/j.neuroimage.2020.117019

Leech, R., Kamourieh, S., Beckmann, C.F., Sharp, D.J., 2011. Fractionating the default mode network: Distinct contributions of the ventral and dorsal posterior cingulate cortex to cognitive control. J. Neurosci. 31, 3217–3224. https://doi.org/10.1523/JNEUROSCI.5626-10.2011

Li, C.-S.R., Yan, P., Bergquist, K.L., Sinha, R., 2007. Greater activation of the “default” brain regions predicts stop signal errors. Neuroimage 38, 640–648. https://doi.org/10.1016/j.neuroimage.2007.07.021

Lieberman, M.D., Straccia, M.A., Meyer, M.L., Du, M., Tan, K.M., 2019. Social, self, (situational), and affective processes in medial prefrontal cortex (MPFC): Causal, multivariate, and reverse inference evidence. Neurosci. Biobehav. Rev. 99, 311–328. https://doi.org/10.1016/j.neubiorev.2018.12.021

Lin, P., Hasson, U., Jovicich, J., Robinson, S., 2011. A neuronal basis for task-negative responses in the human brain. Cereb. Cortex 21, 821–830. https://doi.org/10.1093/cercor/bhq151

Lisofsky, N., Kazzer, P., Heekeren, H.R., Prehn, K., 2014. Investigating socio-cognitive processes in deception: A quantitative meta-analysis of neuroimaging studies. Neuropsychologia 61, 113–122. https://doi.org/10.1016/j.neuropsychologia.2014.06.001

Liu, X., Duyn, J.H., 2013. Time-varying functional network information extracted from brief instances of spontaneous brain activity. Proc. Natl. Acad. Sci. U. S. A. 110, 4392–4397. https://doi.org/10.1073/pnas.1216856110

Lopez-Persem, A., Roumazeilles, L., Folloni, D., Marche, K., Fouragnan, E.F., Khalighinejad, N., Rushworth, M.F.S., Sallet, J., 2020. Differential functional connectivity underlying asymmetric reward-related activity in human and nonhuman primates. Proc. Natl. Acad. Sci. U. S. A. 117, 28452–28462. https://doi.org/10.1073/pnas.2000759117

Mancuso, L., Costa, T., Nani, A., Manuello, J., Liloia, D., Gelmini, G., Panero, M., Duca, S., Cauda, F., 2019. The homotopic connectivity of the functional brain: a meta-analytic approach. Sci. Rep. 9, 3346. https://doi.org/10.1038/s41598-019-40188-3

Mantini, D., Vanduffel, W., 2013. Emerging roles of the brain’s default network. Neuroscientist 19, 76–87. https://doi.org/10.1177/1073858412446202

Mar, R.A., 2011. The neural bases of social cognition and story comprehension. Annu. Rev. Psychol. 62, 103–134. https://doi.org/10.1146/annurev-psych-120709-145406

Marron, T.R., Lerner, Y., Berant, E., Kinreich, S., Shapira-Lichter, I., Hendler, T., Faust, M., 2018. Chain free association, creativity, and the default mode network. Neuropsychologia. https://doi.org/10.1016/j.neuropsychologia.2018.03.018

Mars, R.B., Neubert, F.X., Noonan, M.A.P., Sallet, J., Toni, I., Rushworth, M.F.S., 2012. On the relationship between the "default mode network" and the "social brain." Front. Hum. Neurosci. 1–9. https://doi.org/10.3389/fnhum.2012.00189

Martins, D., Rademacher, L., Gabay, A.S., Taylor, R., Richey, J.A., Smith, D. V, Goerlich, K.S., Nawijn, L., Cremers, H.R., Wilson, R., Bhattacharyya, S., Paloyelis, Y., 2021. Mapping social reward and punishment processing in the human brain: A voxel-based meta-analysis of neuroimaging findings using the Social Incentive Delay task. Neurosci. Biobehav. Rev. https://doi.org/https://doi.org/10.1016/j.neubiorev.2020.12.034

Mason, M.F., Norton, M.I., Horn, John D Van, Wegner, D.M., Grafton, S.T., Macrae, C.N., Mason, M.F., Norton, M.I., Horn, John D Van, Wegner, D.M., Grafton, S.T., Macrae, C.N., Van Horn, J. D., Wegner, D.M., Grafton, S.T., Macrae, C.N., 2007. Wandering Minds: The Default Network and Stimulus-Independent Thought. Science (80-. ). 315, 393–395. https://doi.org/10.1126/science.1131295

Mayseless, N., Eran, A., Shamay-Tsoory, S.G., 2015. Generating original ideas: The neural underpinning of originality. Neuroimage 116, 232–239. https://doi.org/10.1016/j.neuroimage.2015.05.030

Mazoyer, B., Zago, L., Mellet, E., Bricogne, S., Etard, O., Houde, O., Crivello, F., Joliot, M., Petit, L., Tzourio-Mazoyer, N., 2001. Cortical networks for working memory and executive functions sustain the conscious resting state in man. Brain Res Bull 54, 287–298. https://doi.org/10.1016/S0361-9230(00)00437-8

Mennes, M., Kelly, C., Colcombe, S., Xavier Castellanos, F., Milham, M.P., 2013. The extrinsic and intrinsic functional architectures of the human brain are not equivalent. Cereb. Cortex 23, 223–229. https://doi.org/10.1093/cercor/bhs010

Menon, V., Uddin, L.Q., 2010. Saliency, switching, attention and control: a network model of insula function. Brain Struct. Funct. 214, 655–667. https://doi.org/10.1007/s00429-010-0262-0

Mohan, Akansha, Roberto, A.J., Mohan, Abhishek, Lorenzo, A., Jones, K., Carney, M.J., Liogier-Weyback, L., Hwang, S., Lapidus, K.A.B., 2016. The significance of the Default Mode Network (DMN) in neurological and neuropsychiatric disorders: A review. Yale J. Biol. Med. 89, 49–57.

Molnar-Szakacs, I., Uddin, L.Q., 2013. Self-processing and the default mode network: Interactions with the mirror neuron system. Front. Hum. Neurosci. 7, 1–11. https://doi.org/10.3389/fnhum.2013.00571

Müller, V.I., Cieslik, E.C., Laird, A.R., Fox, P.T., Radua, J., Mataix-Cols, D., Tench, C.R., Yarkoni, T., Nichols, T.E., Turkeltaub, P.E., Wager, T.D., Eickhoff, S.B., 2018. Ten simple rules for neuroimaging meta-analysis. Neurosci. Biobehav. Rev. 84, 151–161. https://doi.org/10.1016/j.neubiorev.2017.11.012

Murphy, C., Jefferies, E., Rueschemeyer, S.A., Sormaz, M., Wang, H. ting, Margulies, D.S., Smallwood, J., 2018. Distant from input: Evidence of regions within the default mode network supporting perceptually-decoupled and conceptually-guided cognition. Neuroimage 171, 393–401. https://doi.org/10.1016/j.neuroimage.2018.01.017

Mwilambwe-Tshilobo, L., Spreng, R.N., 2021. Social exclusion reliably engages the default network: A meta-analysis of Cyberball. Neuroimage 227, 117666. https://doi.org/10.1016/j.neuroimage.2020.117666

Najafi, M., McMenamin, B.W., Simon, J.Z., Pessoa, L., 2016. Overlapping communities reveal rich structure in large-scale brain networks during rest and task conditions. Neuroimage 135, 92–106. https://doi.org/10.1016/j.neuroimage.2016.04.054

Newton, A.T., Morgan, V.L., Rogers, B.P., Gore, J.C., 2011. Modulation of steady state functional connectivity in the default mode and working memory networks by cognitive load. Hum. Brain Mapp. 32, 1649–1659. https://doi.org/10.1002/hbm.21138

Ngo, G.H., Eickhoff, S.B., Nguyen, M., Sevinc, G., Fox, P.T., Spreng, R.N., Yeo, B.T.T., 2019. Beyond consensus: Embracing heterogeneity in curated neuroimaging meta-analysis. Neuroimage 200, 142–158. https://doi.org/10.1016/j.neuroimage.2019.06.037

Noonan, K.A., Jefferies, E., Visser, M., Lambon Ralph, M.A., 2013. Going beyond Inferior Prefrontal Involvement in Semantic Control: Evidence for the Additional Contribution of Dorsal Angular Gyrus and Posterior Middle Temporal Cortex. J. Cogn. Neurosci. 25, 1824–1850. https://doi.org/10.1162/jocn_a_00442

Northoff, G., Bermpohl, F., 2004. Cortical midline structures and the self. Trends Cogn. Sci. 8, 102–107. https://doi.org/10.1016/j.tics.2004.01.004

Northoff, G., Heinzel, A., de Greck, M., Bermpohl, F., Dobrowolny, H., Panksepp, J., 2006. Self-referential processing in our brain-A meta-analysis of imaging studies on the self. Neuroimage 31, 440–457. https://doi.org/10.1016/j.neuroimage.2005.12.002

Ochsner, K.N., Beer, J.S., Robertson, E.R., Cooper, J.C., Gabrieli, J.D.E., Kihsltrom, J.F., D'Esposito, M., 2005. The neural correlates of direct and reflected self-knowledge. Neuroimage 28, 797–814. https://doi.org/10.1016/j.neuroimage.2005.06.069

Ochsner, K.N., Knierim, K., Ludlow, D.H., Hanelin, J., Ramachandran, T., Glover, G., Mackey, S.C., 2004. Reflecting upon Feelings: An fMRI Study of Neural Systems Supporting the Attribution of

Emotion to Self and Other. J. Cogn. Neurosci. 16, 1746–1772. https://doi.org/10.1162/0898929042947829

Palla, G., Derényi, I., Farkas, I., Vicsek, T., 2005. Uncovering the overlapping community structure of complex networks in nature and society. Nature 435, 814–818. https://doi.org/10.1038/nature03607

Pedregosa, F., Varoquaux, G., Gramfort, A., Michel, V., Thirion, B., Grisel, O., Blondel, M., Prettenhofer, P., Weiss, R., Dubourg, V., Vanderplas, J., Passos, A., Cournapeau, D., Brucher, M., Perrot, M., Duchesnay, É., 2011. Scikit-learn: Machine Learning in Python. J. Mach. Learn. Res. 12, 2825–2830.

Pessoa, L., 2014. Understanding brain networks and brain organization. Phys. Life Rev. 11, 400–435. https://doi.org/10.1016/j.plrev.2014.03.005

Petersen, S.E., Sporns, O., 2015. Brain Networks and Cognitive Architectures. Neuron 88, 207–219. https://doi.org/10.1016/j.neuron.2015.09.027

Piccoli, T., Valente, G., Linden, D.E.J., Re, M., Esposito, F., Sack, A.T., Salle, F. Di, 2015. The default mode network and the working memory network are not anti-correlated during all phases of a working memory task. PLoS One 10, 1–16. https://doi.org/10.1371/journal.pone.0123354

Prado, J., Weissman, D.H., 2011. Heightened interactions between a key default-mode region and a key task-positive region are linked to suboptimal current performance but to enhanced future performance. Neuroimage 56, 2276–2282. https://doi.org/10.1016/j.neuroimage.2011.03.048

Preti, M.G., Bolton, T.A., Van De Ville, D., 2017. The dynamic functional connectome: State-of-the-art and perspectives. Neuroimage 160, 41–54. https://doi.org/10.1016/j.neuroimage.2016.12.061

Pujol, J., Reixach, J., Harrison, B.J., Timoneda-Gallart, C., Vilanova, J.C., Pérez-Alvarez, F., 2008. Posterior cingulate activation during moral dilemma in adolescents. Hum. Brain Mapp. 29, 910–921. https://doi.org/10.1002/hbm.20436

Raichle, M.E., 2009. A Paradigm Shift in Functional Brain Imaging. J. Neurosci. 29, 12729–12734. https://doi.org/10.1523/JNEUROSCI.4366-09.2009

Raichle, M.E., MacLeod, A.M., Snyder, A.Z., Powers, W.J., Gusnard, D.A., Shulman, G.L., 2001. A default mode of brain function. Proc. Natl. Acad. Sci. 98, 676–682. https://doi.org/10.1073/pnas.98.2.676

Raichle, M.E., Snyder, A.Z., 2007. A default mode of brain function: A brief history of an evolving idea. Neuroimage 37, 1083–1090. https://doi.org/10.1016/j.neuroimage.2007.02.041

Ray, K.L., McKay, D.R., Fox, P.M., Riedel, M.C., Uecker, A.M., Beckmann, C.F., Smith, S.M., Fox, P.T., Laird, A.R., 2013. ICA model order selection of task co-activation networks. Front. Neurosci. 7, 1–12. https://doi.org/10.3389/fnins.2013.00237

Rilling, J.K., Sanfey, A.G., Aronson, J.A., Nystrom, L.E., Cohen, J.D., 2004. The neural correlates of theory of mind within interpersonal interactions. Neuroimage 22, 1694–1703. https://doi.org/10.1016/j.neuroimage.2004.04.015

Ruby, P., Decety, J., 2004. How would You feel versus how do you think She would feel? A neuroimaging study of perspective-taking with social emotions. J. Cogn. Neurosci. 16, 988–999. https://doi.org/10.1162/0898929041502661

Rugg, M.D., Vilberg, K.L., 2013. Brain networks underlying episodic memory retrieval. Curr. Opin. Neurobiol. 23, 255–260. https://doi.org/10.1016/j.conb.2012.11.005

Samartsidis, P., Montagna, S., Laird, A.R., Fox, P.T., Johnson, T.D., Nichols, T.E., 2020. Estimating the prevalence of missing experiments in a neuroimaging meta-analysis. Res. Synth. Methods 11, 866–883. https://doi.org/10.1002/jrsm.1448

Satpute, A.B., Lindquist, K.A., 2019. The Default Mode Network's Role in Discrete Emotion. Trends Cogn. Sci. 23, 851–864. https://doi.org/10.1016/j.tics.2019.07.003

Saxe, R., Kanwisher, N., 2003. People thinking about thinking people: The role of the temporo-parietal junction in "theory of mind." Neuroimage 19, 1835–1842. https://doi.org/10.1016/S1053-8119(03)00230-1

Saxe, R., Powell, L.J., 2006. It's the Thought That Counts. Psychol. Sci. 17, 692–699. https://doi.org/10.1111/j.1467-9280.2006.01768.x

Schacter, D.L., Addis, D.R., Buckner, R.L., 2008. Episodic simulation of future events: Concepts, data, and applications. Ann. N. Y. Acad. Sci. 1124, 39–60. https://doi.org/10.1196/annals.1440.001

Schacter, D.L., Addis, D.R., Buckner, R.L., 2007. Remembering the past to imagine the future: The prospective brain. Nat. Rev. Neurosci. 8, 657–661. https://doi.org/10.1038/nrn2213

Schilbach, L., Bzdok, D., Timmermans, B., Fox, P.T., Laird, A.R., Vogeley, K., Eickhoff, S.B., 2012. Introspective Minds: Using ALE meta-analyses to study commonalities in the neural correlates of emotional processing, social & unconstrained cognition. PLoS One 7. https://doi.org/10.1371/journal.pone.0030920

Schneider, B., Koenigs, M., 2017. Human lesion studies of ventromedial prefrontal cortex. Neuropsychologia 107, 84–93. https://doi.org/10.1016/j.neuropsychologia.2017.09.035

Schultz, W., 2015. Neuronal reward and decision signals: From theories to data. Physiol. Rev. 95, 853–951. https://doi.org/10.1152/physrev.00023.2014

Seli, P., Risko, E.F., Smilek, D., Schacter, D.L., 2016. Mind-Wandering With and Without Intention. Trends Cogn. Sci. 20, 605–617. https://doi.org/10.1016/j.tics.2016.05.010

Sha, Z., Wager, T.D., Mechelli, A., He, Y., 2019. Common Dysfunction of Large-Scale Neurocognitive Networks Across Psychiatric Disorders. Biol. Psychiatry 85, 379–388. https://doi.org/10.1016/j.biopsych.2018.11.011

Sha, Z., Xia, M., Lin, Q., Cao, M., Tang, Y., Xu, K., Song, H., Wang, Z., Wang, F., Fox, P.T., Evans, A.C., He, Y., 2018. Meta-Connectomic Analysis Reveals Commonly Disrupted Functional Architectures in Network Modules and Connectors across Brain Disorders. Cereb. Cortex 28, 4179–4194. https://doi.org/10.1093/cercor/bhx273

Shine, J.M., Breakspear, M., Bell, P.T., Ehgoetz Martens, K., Shine, R., Koyejo, O., Sporns, O., Poldrack, R.A., 2019. Human cognition involves the dynamic integration of neural activity and neuromodulatory systems. Nat. Neurosci. 22, 289–296. https://doi.org/10.1038/s41593-018-0312-0

Shirer, W.R., Ryali, S., Rykhlevskaia, E., Menon, V., Greicius, M.D., 2012. Decoding Subject-Driven Cognitive States with Whole-Brain Connectivity Patterns. Cereb. Cortex 22, 158–165. https://doi.org/10.1093/cercor/bhr099

Shulman, G.L., Fiez, J.A., Corbetta, M., Buckner, R.L., Miezin, F.M., Raichle, M.E., Petersen, S.E., 1997. Common Blood Flow Changes across Visual Tasks: II. Decreases in Cerebral Cortex. J. Cogn. Neurosci. 9, 648–663. https://doi.org/10.1162/jocn.1997.9.5.648

Smallwood, J., McSpadden, M., Schooler, J.W., 2008. When attention matters: The curious incident of the wandering mind. Mem. Cogn. 36, 1144–1150. https://doi.org/10.3758/MC.36.6.1144

Smith, S.M., Fox, P.M.T.M.T., Miller, K.L., Glahn, D.C., Fox, P.M.T.M.T., Mackay, C.E., Filippini, N., Watkins, K.E., Toro, R., Laird, A.R., Beckmann, C.F., 2009. Correspondence of the brain's functional architecture during activation and rest. Proc. Natl. Acad. Sci. 106, 13040–13045. https://doi.org/10.1073/pnas.0905267106

Sonuga-Barke, E.J.S., Castellanos, F.X., 2007. Spontaneous attentional fluctuations in impaired states and pathological conditions: A neurobiological hypothesis. Neurosci. Biobehav. Rev. 31, 977–986. https://doi.org/10.1016/j.neubiorev.2007.02.005

Sormaz, M., Murphy, C., Wang, H.T., Hymers, M., Karapanagiotidis, T., Poerio, G., Margulies, D.S., Jefferies, E., Smallwood, J., 2018. Default mode network can support the level of detail in experience during active task states. Proc. Natl. Acad. Sci. U. S. A. 115, 9318–9323. https://doi.org/10.1073/pnas.1721259115

Spiers, H.J., Maguire, E.A., 2006. Spontaneous mentalizing during an interactive real world task: An fMRI study. Neuropsychologia 44, 1674–1682. https://doi.org/10.1016/j.neuropsychologia.2006.03.028

Sporns, O., Tononi, G., Kötter, R., 2005. The human connectome: A structural description of the human brain. PLoS Comput. Biol. 1, 0245–0251. https://doi.org/10.1371/journal.pcbi.0010042

Spreng, R.N., 2012. The Fallacy of a “Task-Negative” Network. Front. Psychol. 3, 1–5. https://doi.org/10.3389/fpsyg.2012.00145

Spreng, R.N., Andrews-Hanna, J.R., 2015. The Default Network and Social Cognition. Brain Mapp. An Encycl. Ref. 3, 165–169. https://doi.org/10.1016/B978-0-12-397025-1.00173-1

Spreng, R.N., DuPre, E., Selarka, D., Garcia, J., Gojkovic, S., Mildner, J., Luh, W.-M., Turner, G.R., 2014. Goal-Congruent Default Network Activity Facilitates Cognitive Control. J. Neurosci. 34, 14108–14114. https://doi.org/10.1523/JNEUROSCI.2815-14.2014

Spreng, R.N., Gerlach, K.D., Turner, G.R., Schacter, D.L., 2015. Autobiographical Planning and the Brain: Activation and Its Modulation by Qualitative Features. J. Cogn. Neurosci. 27, 2147–2157. https://doi.org/10.1162/jocn_a_00846

Spreng, R.N., Grady, C.L., 2010. Patterns of brain activity supporting autobiographical memory, prospection, and theory of mind, and their relationship to the default mode network. J. Cogn. Neurosci. 22, 1112–1123. https://doi.org/10.1162/jocn.2009.21282

Spreng, R.N., Mar, R.A., Kim, A.S.N., 2009. The Common Neural Basis of Autobiographical Memory, Prospection, Navigation, Theory of Mind, and the Default Mode: A Quantitative Meta-analysis. J. Cogn. Neurosci. 21, 489–510. https://doi.org/10.1162/jocn.2008.21029

Spreng, R.N., Sepulcre, J., Turner, G.R., Stevens, W.D., Schacter, D.L., 2013. Intrinsic Architecture Underlying the Relations among the Default, Dorsal Attention, and Frontoparietal Control Networks of the Human Brain. J. Cogn. Neurosci. 25, 74–86. https://doi.org/10.1162/jocn_a_00281

Spreng, R.N., Stevens, W.D., Chamberlain, J.P., Gilmore, A.W., Schacter, D.L., 2010. Default network activity, coupled with the frontoparietal control network, supports goal-directed cognition. Neuroimage 53, 303–317. https://doi.org/10.1016/j.neuroimage.2010.06.016

Sridharan, D., Levitin, D.J., Menon, V., 2008. A critical role for the right fronto-insular cortex in switching between central-executive and default-mode networks. Proc. Natl. Acad. Sci. 105, 12569–12574. https://doi.org/10.1073/pnas.0800005105

Svoboda, E., McKinnon, M.C., Levine, B., 2006. The functional neuroanatomy of autobiographical memory: A meta-analysis. Neuropsychologia 44, 2189–2208. https://doi.org/10.1016/j.neuropsychologia.2006.05.023

Toro-Serey, C., Tobyne, S.M., McGuire, J.T., 2020. Spectral partitioning identifies individual heterogeneity in the functional network topography of ventral and anterior medial prefrontal cortex. Neuroimage 205, 116305. https://doi.org/10.1016/j.neuroimage.2019.116305

Toro, R., Fox, P.T., Paus, T., 2008. Functional coactivation map of the human brain. Cereb. Cortex 18, 2553–2559. https://doi.org/10.1093/cercor/bhn014

Turkeltaub, P.E., Eden, G.F., Jones, K.M., Zeffiro, T.A., 2002. Meta-analysis of the functional neuroanatomy of single-word reading: Method and validation. Neuroimage 16, 765–780. https://doi.org/10.1006/nimg.2002.1131

Turkeltaub, P.E., Eickhoff, S.B., Laird, A.R., Fox, M., Wiener, M., Fox, P., 2012. Minimizing within-experiment and within-group effects in activation likelihood estimation meta-analyses. Hum. Brain Mapp. 33, 1–13. https://doi.org/10.1002/hbm.21186

Uddin, L.Q., Clare Kelly, A.M., Biswal, B.B., Xavier Castellanos, F., Milham, M.P., 2009. Functional connectivity of default mode network components: Correlation, anticorrelation, and causality. Hum. Brain Mapp. 30, 625–637. https://doi.org/10.1002/hbm.20531

Uddin, L.Q., Iacoboni, M., Lange, C., Keenan, J.P., 2007. The self and social cognition: the role of cortical midline structures and mirror neurons. Trends Cogn. Sci. 11, 153–157. https://doi.org/10.1016/j.tics.2007.01.001

Uddin, L.Q., Yeo, B.T.T., Spreng, R.N., 2019. Towards a Universal Taxonomy of Macro-scale Functional Human Brain Networks. Brain Topogr. 32, 926–942. https://doi.org/10.1007/s10548-019-00744-6

Vatansever, Deniz, Menon, D.K., Manktelow, A.E., Sahakian, B.J., Stamatakis, E.A., 2015. Default mode dynamics for global functional integration. J. Neurosci. 35, 15254–15262. https://doi.org/10.1523/JNEUROSCI.2135-15.2015

Vatansever, D., Menon, D.K., Manktelow, A.E., Sahakian, B.J., Stamatakis, E.A., 2015. Default mode network connectivity during task execution. Neuroimage 122, 96–104. https://doi.org/10.1016/j.neuroimage.2015.07.053

Vatansever, D., Menon, D.K., Stamatakis, E.A., 2017. Default mode contributions to automated information processing. Proc. Natl. Acad. Sci. U. S. A. 114, 12821–12826. https://doi.org/10.1073/pnas.1710521114

Verdejo-García, A., Bechara, A., 2009. A somatic marker theory of addiction. Neuropharmacology 56, 48–62. https://doi.org/10.1016/j.neuropharm.2008.07.035

Voytek, B., Knight, R.T., 2015. Dynamic network communication as a unifying neural basis for cognition, development, aging, and disease. Biol. Psychiatry 77, 1089–1097. https://doi.org/10.1016/j.biopsych.2015.04.016

Wang, S., Tepfer, L.J., Taren, A.A., Smith, D. V., 2020. Functional parcellation of the default mode network: a large-scale meta-analysis. Sci. Rep. 10. https://doi.org/10.1038/s41598-020-72317-8

Weissman, D.H., Roberts, K.C., Visscher, K.M., Woldorff, M.G., 2006. The neural bases of momentary lapses in attention. Nat. Neurosci. 9, 971–978. https://doi.org/10.1038/nn1727

Wen, T., Mitchell, D.J., Duncan, J., 2020. The Functional Convergence and Heterogeneity of Social, Episodic, and Self-Referential Thought in the Default Mode Network. Cereb. Cortex. https://doi.org/10.1093/cercor/bhaa166

Xue, G., Lu, Z., Levin, I.P., Weller, J.A., Li, X., Bechara, A., 2009. Functional dissociations of risk and reward processing in the medial prefrontal cortex. Cereb. Cortex 19, 1019–1027. https://doi.org/10.1093/cercor/bhn147

Yang, X.F., Bossmann, J., Schiffhauer, B., Jordan, M., Immordino-Yang, M.H., 2013. Intrinsic default mode network connectivity predicts spontaneous verbal descriptions of autobiographical memories during social processing. Front. Psychol. 3, 1–10. https://doi.org/10.3389/fpsyg.2012.00592

Yeo, B.T.T., Krienen, F.M., Chee, M.W.L., Buckner, R.L., 2014. Estimates of segregation and overlap of functional connectivity networks in the human cerebral cortex. Neuroimage 88, 212–227. https://doi.org/10.1016/j.neuroimage.2013.10.046

Yeo, B.T.T., Krienen, F.M., Sepulcre, J., Sabuncu, M.R., Lashkari, D., Hollinshead, M., Roffman, J.L., Smoller, J.W., Zollei, L., Polimeni, J.R., Fischl, B., Liu, H., Buckner, R.L., Thomas Yeo, B.T., Krienen, F.M., Sepulcre, J., Sabuncu, M.R., Lashkari, D., Hollinshead, M., Roffman, J.L., Smoller, J.W., Zollei, L., Polimeni, J.R., Fischl, B., Liu, H., Buckner, R.L., 2011. The organization of the human cerebral cortex estimated by intrinsic functional connectivity. J. Neurophysiol. 106, 1125–1165. https://doi.org/10.1152/jn.00338.2011
